# Deep physico-chemical characterization of the serum antibody response using a dual-titration microspot assay

**DOI:** 10.1101/2023.03.14.532012

**Authors:** Ágnes Kovács, Zoltán Hérincs, Krisztián Papp, Jakub Zbigniew Kaczmarek, Daniel Nyberg Larsen, Pernille Stage, László Bereczki, Eszter Ujhelyi, Tamás Pfeil, József Prechl

## Abstract

Antigen specific humoral immunity can be characterized by the analysis of serum antibodies. While serological assays for the measurement of specific antibody levels are available, these are not quantitative in the biochemical sense. Yet, understanding humoral immune responses quantitatively on the systemic level would need a universal, complete, quantitative, comparable measurement method of antigen specific serum antibodies of selected immunoglobulin classes. Here we describe a fluorescent, dual-titration immunoassay, which provides the physico-chemical parameters that are both necessary and sufficient to quantitatively characterize the humoral immune response. We define the theory of the approach that is based on physical chemistry. For validation of theory, we used recombinant receptor binding domain of SARS-CoV-2 as antigen on microspot arrays and varied the concentration of both the antigen and serum antibodies from infected persons to obtain a measurement matrix of binding data. Both titration curves were simultaneously fitted using an algorithm based on the generalized logistic function and adapted for analyzing thermodynamic variables of binding. We obtained equilibrium affinity constants and chemical potentials for distinct antibody classes. These variables reflect the quality and the effective quantity of serum antibodies, respectively. The proposed fluorescent dual-titration microspot immunoassay can generate truly quantitative serological data that is suitable for immunological, medical and systems biological analysis.

## 1. Introduction

The adaptive immune system maintains host integrity by controlling the levels of molecules and cells over a very wide range. The humoral adaptive immune system achieves this by employing effector molecules with tunable specificity and efficiency. These molecules, antibodies (Ab), evolve with the help of B lymphocytes that use them both as sensors for their survival and as effectors for the removal of target molecules. To protect against viral invasion of host cells, antibodies with enhanced effector activity (1) are produced and persist over time (2). However, the emergence of novel viruses with enhanced ability to enter and spread in the host may overwhelm before such responses might take place (3). Since antibodies circulating in blood are good indicators of the status of humoral immunity, the characterization of serum antibodies, Ab serology, plays a critical role in immunodiagnostics.

Currently two main approaches dominate serological methods of antigen (Ag) specific Ab measurements: titration and single-point assays. Titration is the use of a series of gradually diluted serum and the determination of mid-point or end-point titer from the dilution curve. This approach does incorporate the measurement of effects due to changing reaction conditions, but the readout most often neglects curve shape and focuses on a single unitless parameter: the titer. Single-point measurements are optimized for diagnostic power (sensitivity and specificity) (4–6) and neglect potential effects of serum dilution. The readout of single-point measurements is an arbitrary though standardized unit of reactivity. Virus neutralization (7) and surrogate neutralization assays (8,9) can also be carried out in the single-point or titration format. What is common in these assays is that none of them varies or takes into account the density of the target molecule used in the assay: the solid-phase coupled Ag or the virus receptor on cells.

Interactions between Ab and Ag constitute the basis of immunoassays and while several physical and mathematical models have been developed and utilized to characterize them, yet these are primarily assays wherein the analyte is the Ag and determination of Ag concentration is the aim. Less attention has been paid to the development of models dedicated to the special requirements of circulating serum Ab detection, that is, when the analyte is not a single molecular species but the collection of serum Ab. Serum Abs are highly heterogeneous with respect to epitope recognition, affinity, concentrations and structure. A blood serum sample in fact reflects immunological memory of past infections, diseases, along with current immunological activity. While serological assay results are given in units of activity, referring to binding or biochemical activity, no clear relationship to thermodynamic activity has been established (10). Early approaches of Ag-Ab interaction modeling, based on predictions of molecular properties (11), suggested that too many physical properties of the interacting molecules should be estimated.

The difficulty of serum Ab quantification lies in the fact that we are dealing with two unknown variables: average effective affinities and concentrations of Abs that bind to the Ag in the assay. Any approach aiming to quantify serum Abs should consider estimating both. Traditionally equilibrium dialysis was the method of choice for measuring affinity constants in solution when radioimmunoassays were used. These assays used a constant concentration of Ab (e.g., 1:100 dilution of serum) and varying concentrations of Ag, calculating the concentration of free radioactively labeled Ag, once equilibrium was established, from the radioactivity. The Scatchard plot was used for calculating the median and mean equilibrium dissociation constant (KD) (12) and for the assessment of affinity heterogeneity and the Sips plot was used for quantifying heterogeneity (13,14). Current approaches employ novel technologies for assessing affinity and concentration. Lippok et al. used microscale thermopheresis to measure both affinity and concentration of polyclonal Abs in solution (15). Fiedler et al. utilized diffusional sizing of labeled Ag in a microfluidic device for KD and concentration estimation (16). Tang et al. described the combination of ELISA and quantitative mass spectrometry, an approach that allows quantitation of bound Ab (17). These novel approaches generate results in universal biochemical units but still fall short of revealing important biochemical properties of serum Abs of selected classes.

Microspot immunoassays represent special measurement conditions with respect to the relative amounts of the reactants: the mass of the reactant in the microspot is negligible to the mass of the reactant in solution. This property is exploited in ambient analyte immunoassays (18), where capture Abs are printed as microspots and are used for determining the concentration of Ag in solution. Because of the negligible concentration change caused by the capture of Ag from solution, this setup is ideal for concentration measurements (19). Microspotted Ag can be used for the measurement of serum Ab binding, with the assumption that captured Ab causes negligible change in the composition of the solution (20). From the physico-chemical point of view, these conditions correspond to measurements carried out at infinite Ag dilution, allowing the estimation of limiting thermodynamic parameters of the binding reaction.

As a model system for our proof-of-concept study we chose to measure the serum Ab response against a domain of the spike protein of the pandemic coronavirus. Infection with SARS-CoV-2 results in the appearance of IgM (21), IgG (22–24) and IgA (25) directed against various viral components (26–29). The response builds on immunological imprints from common cold viruses (30), is connected to disease severity (31–35) and lasts several months (36,37). Vaccination also induces Ab production, therefore serological measurements can help assess vaccine immunogenicity and estimate protectivity (38–41), durability (42) and cross-reactivity with emerging new variants (43). The various indicators of humoral immune response, such as titers (44,45), avidity (46), glycosylation (33) of virus specific Abs are all related to clinical aspects of the infection and disease, but these relationships are highly variable, not definite (3), partially because of the difficulties in comparing different measurement methods (32).

We recently developed a microspot-based, dual-titration immunoassay for the estimation of affinity distribution for multiple immunoglobulin classes (20) utilizing an advanced fitting algorithm (47). Here we describe a further improvement of the assay and demonstrate its applicability for the characterization of anti-SARS-CoV-2 Ab response, by the estimation of chemical thermodynamic variables of SARS-CoV-2 RBD specific Abs of various classes. This approach not only produces universal biochemical units of measurement for the Ab isotype of interest but at the same time also provides insight into the chemical thermodynamics of a complex system. While natural logarithm of the data was used for fitting (abbreviated with ln), concentrations and KD values are displayed as base-10 logarithmic values (abbreviated with log) for easier interpretation.

## 2. Materials and Methods

### 2.1 Serum samples

Commercially available serum from confirmed COVID-19 positive and negative subjects with available IgG and IgM test results (RayBiotech CoV-PosSet-2) was obtained from THP Medical Products Vertriebs GmbH, Vienna. COVID-19 positive samples were tested by the manufacturer using their COVID-19 ELISA kits for IgG and IgM measurements (RayBiotech, Georgia, USA).

#### 2.2.1 Gene synthesis, protein expression and characterization of SARS-CoV-2 Spike RBD 319-541

SARS-CoV-2 Spike RBD 319-541 sequence was based on the first strain of SARS-CoV-2 isolated from a clinical patient on January the 6th 2020, (GISAID: EPI_ISL_402119). The sequence was synthesized to include the Tyr-Pho signal peptide and a N-terminal hexahistidine (6xHIS) tag. The construct was synthesized and cloned commercially into pcDNA3.4-TOPO plasmid (Life Technologies B.V. Europe). SARS-CoV-2 Spike RBD 319-541 recombinant protein (OBA0101, Ovodan Biotech) was expressed in 25 mL culture using the Expi293F expression system (#A14635; ThermoFisher Scientific) according to manufacturer’s instructions. Expressed proteins were harvested by centrifugation 6 days post transfection and immediately purified from the supernatant on a Ni-NTA Superflow column (#30430, Qiagen). Eluted protein fraction was buffer exchanged into phosphate-buffered saline solution pH 7.4 using a HiPrep 26/10 Desalting column (#GE17-5087-01, Cytiva) and stored at -20°C.

The obtained protein fraction was subjected to Sodium Dodecyl Sulfate Polyacrylamide gel electrophoresis (SDS-PAGE) using RunBlue (#ab270467 Abcam) 4-12% Bis-Tris polyacrylamide gel. Prior to loading, the samples have been mixed with 2,5µl NuPAGE LDS (4x) sample buffer (Life Technologies) each and incubated in 70 °C for 10 minutes in a glass container filled with water, heated on a VMS-C7 heating block (VWR). Afterwards the samples have been briefly centrifuged using MiniSpin Plus Centrifuge at 1000 RPM for 15 seconds (Eppendorf). Electrophoresis was performed by using XCell SureLock Electrophoresis Cell (Novex) and Easy Power 500 (Invitrogen) in a non-reduced environment for 45 minutes at 200V and 110mA. Electrophoresis was carried out in a 1X MOPS-SDS Buffer (VWR). Staining was performed by Coomassie Simply Blue Safe Stain (#LC6060 Invitrogen) for 1 hour on a PS-M3D orbital shaker (Grand Bio). The gel was destained in deionized water overnight and visualized by using Bio-Rad Gel Doc XR – Molecular Imager. The molecular weight marker used was peq Marker Gold V (VWR). The detailed protocol and PAGE image are available as Supplementary data.

#### 2.2.2 SARS-CoV-2 recombinant RBD sequence characterization by Mass Spectrometry

A total of 20 µg of SARS-CoV-2 Spike RBD 319-541 was reduced with dithiothreitol (20 mM) for 30 minutes at 57 degrees and alkylated with iodoacetamide (54 mM) for 20 minutes at room temperature and in the dark, the reaction was stopped with dithiothreitol. The reduced and alkylated SARS-CoV-2 Spike RBD 319-541 was split into two batches, where the first was treated with 2% homemade methylated trypsin [1] for an hour at 57 degrees and PNGaseF (0.5 µl) (Promega, V4831) for an hour at 37 degrees, while the other was only treated with 2% trypsin for an hour at 57 degrees.

The two batches were micro-purified prior to analysis by mass spectrometry. The micro-purification was performed in accordance with Rappsilber *et al*. (48), where a p200 tip was plugged with M3 material [Empore octyl C8, 66882-U] and 1 μl of R2 material [Poros 20 R2 Applied Biosystems, Part no. 1-1129-06] was added. The stage tip was then activated using 100% acetonitrile [VWR, 83640.290], followed by equilibration with 0.1% TFA [MERCK, 200-929-3], sample was then 1 μg of sample was loaded into 40 μl of 0.1% TFA, followed by washing with 0,1% TFA. The sample was then eluted from the stage tip using first 50% acetonitrile, 0.1% TFA and secondly using 70% acetonitrile, 0.1% TFA. The eluted samples were then lyophilized using an Eppendorf vacuum centrifuge [VWR, 20318.297], prior to running the samples were resuspended in 6 μl 0.1% formic acid [maker needed]. Both batches were run in duplicate, using 1 μg of sample per run, with a standard liquid chromatographic tandem mass spectrometric analysis on an Orbitrap Exploris™ 480 Mass Spectrometer from Thermo Fisher Scientific. The follow main settings were used; MS1 resolution: 120000, scan range (m/z): 350-1400, included charge state(s): 1.0e4, dynamic exclusion after 1 time with an exclusion time of 30 seconds, MS2 resolution: 30000, isolation window MS2 (m/z): 0.8, first mass MS2 (m/z): 110, data type: centroid.

The data files from the mass spectrometer were converted using MSConvert from ProteoWizard [https://proteowizard.sourceforge.io/], followed by data search and analysis in GPMAW from Lighthouse Data [http://www.gpmaw.com/].

The mass spectrometry proteomics data have been deposited to the ProteomeXchange Consortium via the PRIDE (45) partner repository with the dataset identifier PXD040415. Further details of protein characterization are available online as Supplementary Data.

### 2.3 Dual-titration microspot immunoassay

Maps of the layout of slides and subarrays, along with a detailed description of the protocol are available online as Supplementary data.

#### 2.3.1 Microarray production

Experiments were carried out on hydrogel-coated glass slides (Nexterion Slide H, Schott Minifab, Jena) by using a BioOdyssey Calligrapher MiniArrayer (BioRad). A 14-point dilution series of RBD was prepared with a combination of a ½ and ⅓ diluting series, and spotted on slides in triplicates. The final concentration gradient steps were: 16.66, 8.33, 5.55, 4.16, 2.08, 1.85, 1.04, 0.61, 0.52, 0.26, 0.20, 0.13, 0.068 and 0.065 µM. Slides were dried for 1h at 37°C then soaked in 0.1 M Tris buffer (pH=8.0) for 1h at 37°C in order to block reactive residues on the surface. Once prepared, slides were kept in sealed non-transparent bags at 4°C.

#### 2.3.2 Sample handling and signal detection

Dried arrays were rehydrated in 110 µl PBS (3×5 minutes) before using, then sub-arrays were incubated in 70 µl diluted sample at 37 °C for 1 hour. Sample dilution was carried out in PBS-BSA-Tween (PBS, 0.5% BSA, 0.05% Tween 20). Serum treated slides were washed in 0.05% Tween-PBS, then incubated at room temperature for 30 minutes with fluorescently labeled Abs that were diluted in the blocking buffer (0.05% Tween 20, 2% BSA, PBS). The first mix of detecting Abs was composed of the following: anti-human-IgG F(ab’)_2_ – Alexa488 (Jackson, Ref.:109-646-097), anti-human-IgA – Alexa647 (Jackson, Ref.:109-606-011), anti-human-IgM – Cy3 (Jackson, Ref.: 109-166-129). Chips were washed again and following drying, slides were scanned using SensoSpot fluorescent microarray scanner (Sensovation AG, Stockach, Germany). Fluorescence signals below ln(FI)=4 were excluded from further analysis.

#### 2.3.3 Analysis of the microarray data

Images of the slides were analyzed with GenePix Pro 6.0 software after visual inspection. Spots were recognized, aligned and analyzed by the program, then gpr files containing the spot data were created. Relative fluorescence intensity (RFI) values were calculated for each spot using the feature’s median RFI value of which the feature’s local background was subtracted individually for each feature. Further analysis was carried out by using the statistical programming environment R (version 3.5.2).

### 2.4 Curve fitting

The general theoretical approach to data analysis was similar to that described recently (20). We use a linear model for polyclonal reactions (49) taking into consideration that a bound Ab inhibits nearby free Ags from forming complexes with other Abs such that the concentration of immune complexes is a logistic function of the logarithm of the total Ag concentration. Since the logarithm of fluorescent intensity is proportional to the logarithm of bound Ab concentration, we assume that the fluorescent intensity of detected Abs is a Richards function (R) of the logarithm of total Ag concentration (see Supplementary text 1 of (20)), with the following parametrization:

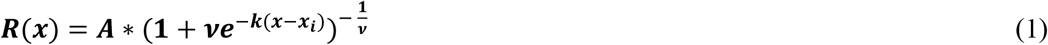

with k = 1, where x is logarithm of Ag concentration ln[Ag], A is the signal corresponding to total Ab concentration [Ab] (limit of function R(x) at infinity), x_i_ is the inflection point of function R, ν is the asymmetry parameter.

The upper limit of the fluorescence, A, depends on the dilution of serum: the less diluted the serum sample, the higher the Ab concentration. We used the reciprocal of serum dilution as a surrogate Ab concentration. Ten times diluted serum corresponds to a concentration of 0.1. The fluorescent signal intensity of bound Ab is thus determined by both Ag and Ab concentrations, which we express using the logarithm total Ag concentration, x=ln[Ag], and the logarithm of the reciprocal of serum dilution factor, y=ln[Ab], as the product of two generalized logistic functions (R_1_(x)*R_2_(y)) with k=1 in the form

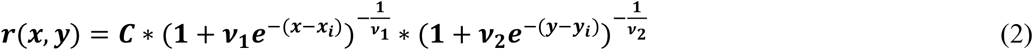

where C is the fluorescent signal corresponding to the maximal concentration of AbAg complexes. Both terms can be normalized to their own inflection points

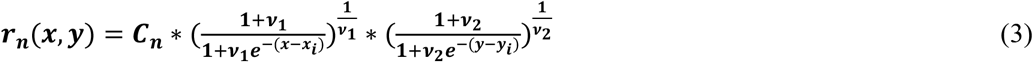

with the visual advantage of moving x_i_ and y_i_ to the same horizontal line z=1 and with the theoretical advantage of introducing relative thermodynamic activity (see explanation later). In this form C_n_ corresponds to the fluorescent signal of AbAg complexes at the inflection point of the functions.

A logarithmic transformation converts the proportional variance pattern to a constant variance pattern and thus the conversion makes the transformed data more suitable for fitting the model. The above multiplicative relationship then changes to an additive one (lnR_1_(x)+lnR_2_(y)) in the form of

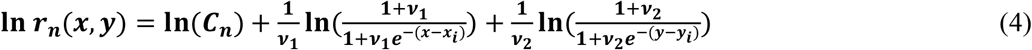

In order to reduce the number of estimated variables and to introduce a variable common to both titration curves, we used the relationship between the asymmetry parameters of Ab and Ag titration curves (Prechl 2024)

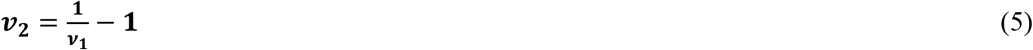

This normalized, generalized logistic model on the log-log scale was fit to the data, and parameter estimates and 95% confidence intervals were obtained using the R software (version 3.5.2). Nonlinear least squares estimates for the model parameters were calculated using the Gauss-Newton algorithm of the *nls* function from the statistical software package R (version 3.5.2). Confidence intervals generated for the model parameters were based on the Wald-based technique.

The concentration of AbAg complexes is a cumulative measure of the activity of both components: by increasing the concentration of either, more and more complexes are formed. Activity, on the other hand, is the rate of change of complex formation, which is given by the partial derivatives of the fitted functions. Once the above parameters of the functions are estimated, the relative thermodynamic activity of the complex at various Ab and Ag concentrations can be obtained from the products of the derivatives of Richards functions R_1_’(ln[Ag])*R_2_’(ln[Ab]) as shown in Figure 1.

**Figure 1.**
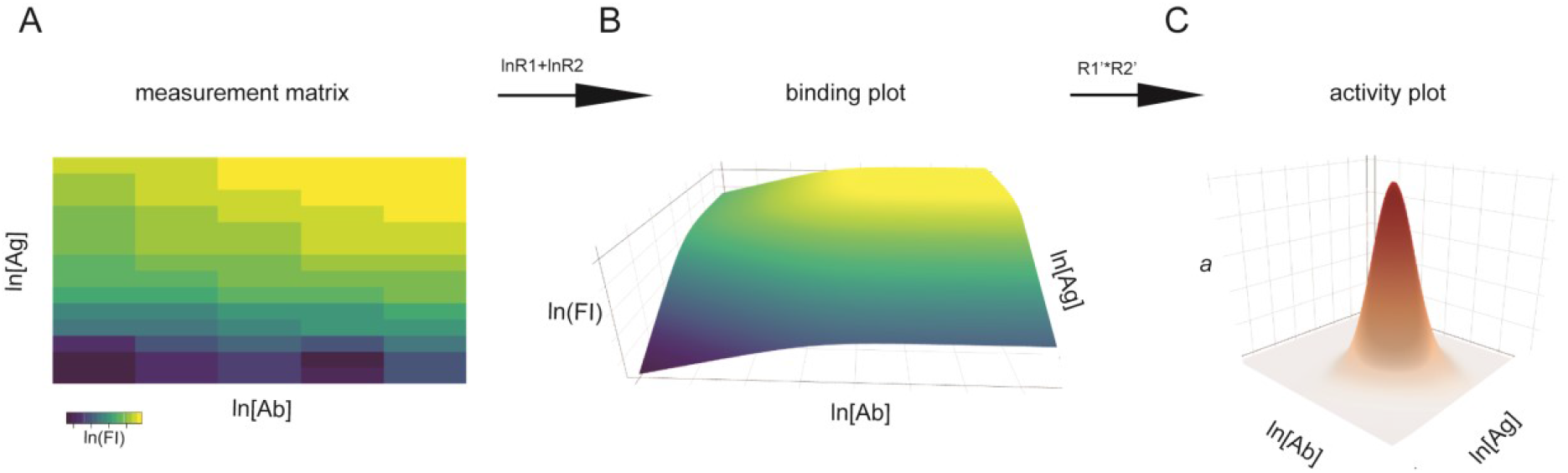
Calculation of standard state of AbAg complexes. A) A measurement matrix is created from the dual titration, using the logarithm of Ab and Ag concentrations (ln[Ab], ln[Ag]) and of the detected fluorescent intensity signals (ln(FI)). B) The combination of lnR functions (lnR1+lnR2, where 1 and 2 stand for ln[Ag] and ln[Ab], respectively) is used for fitting binding data, which corresponds to obtaining the parameters of an AbAg binding surface in three dimensions. C) An activity plot obtained from the product of the derivatives of two Richards functions (R1’*R2’), identifies the standard state as the condition with highest relative thermodynamic activity of AbAg complexes.

**Figure 2.**
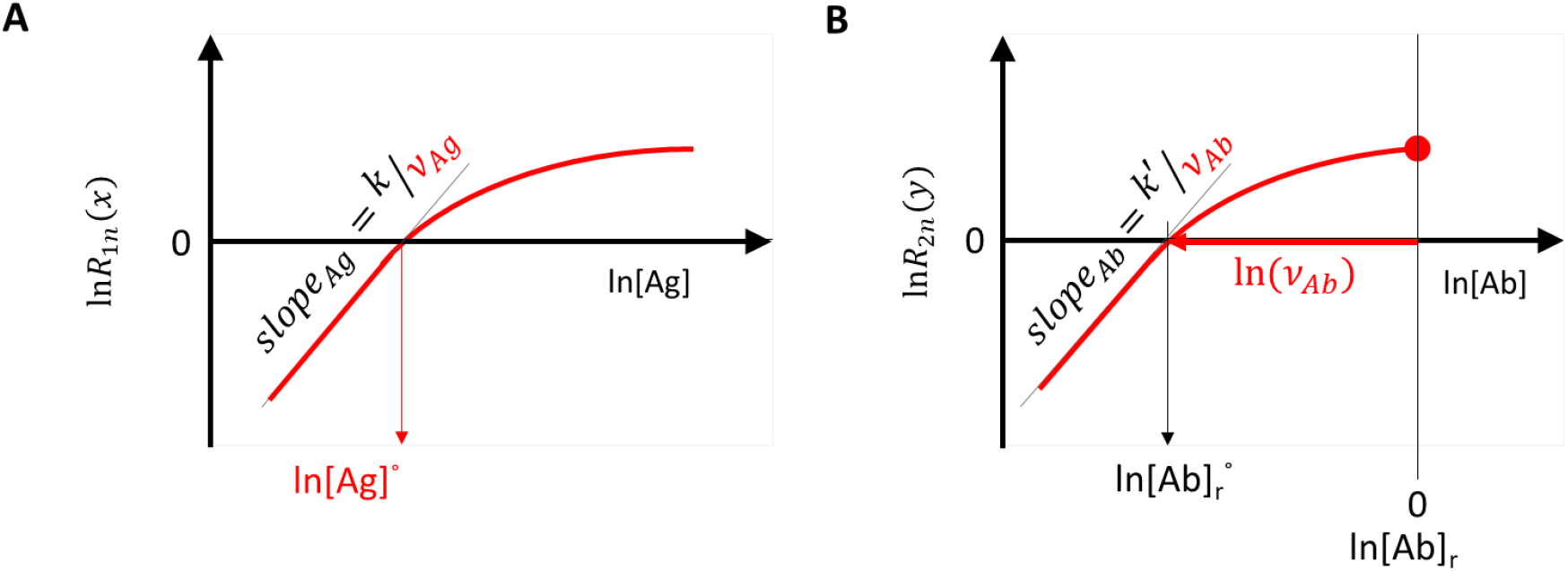
Interpretation of fitted binding curves. The binding curves obtained by titration of A) Ag and B) Ab are shown. A) The logarithm of normalized R1 function intersects the horizontal axis at the standard Ag concentration. B) The distance of the intersection of logarithm of normalized R2 function from the origo (red arrow) is the logarithm of Ab concentration relative to standard Ab concentration. The slope of the left asymptotes is determined by the respective asymmetry parameters. [Ag]: concentration of printed Ag; [Ab]_r_: relative Ab concentration, 1/serum dilution factor

### 2.5 Statistical analysis of results

Throughout the paper we use ln for natural logarithm and log for base 10 logarithm. While curve fitting was carried out on ln transformed data, for the visualization and comparative analysis of results we use log data as is conventional in immunochemistry. Linear regression was used to obtain equations for the calibration of fluorescence and Ab concentrations of IgA, IgG and IgM. Random effects models were used to analyze ν, log(KD), log(KD’_Ag_), log[AbAg]°, and μ_Ab_, with group (positive, negative), antibody class (IgA, IgG, and IgM) and their interaction as fixed effects, and with patient ID as a random effect. Tukey-adjusted p-values were calculated for multiple comparisons between the two groups at each level of anybody class. The statistical analysis was performed using the glht() function from the R multcomp package (R version 4.3.2).

## 3. Results

### 3.1 Definition of standard reference state of polyclonal Ab interactions

In chemical thermodynamics the standard state is a precisely defined reference state, which serves as a common reference point for the comparison of thermodynamic properties. The standard chemical potential is arbitrarily defined in any system in a way to suit the description of the system. For the purpose of chemical reactions pressure, temperature and material composition can define a standard state. For the purpose of an immunoassay composition is critical: the chemical potential of an Ab solution, besides the affinity and concentration of the Ab itself, is determined by the quality and the concentration of the target molecules, Ag. In our model Ab binding to Ag microspots is described by the product of two Richards functions (20). Each of the Richards functions represents the growth of the relative concentration of bound reactants, one as a function of the logarithm of Ag density, the other as a function of the logarithm of Ab concentration (Fig. 1A). The product of these functions can be visualized by a 3D surface plot (Fig. 1B). In this system the inflection points of the two functions can serve as the origin of coordinates. We therefore define the standard reference state of a particular serum Ab and particular Ag mixture as the concentrations of Ab and Ag required to reach the point of inflection along both titration axes and the concentration of AbAg complex at the intersection of inflection points. In general, it means that each serum Ab-Ag pair will have a distinct standard state and the identified standard reference state concentrations will therefore characterize the serum sample in terms of its thermodynamic activity against the tested Ag. We shall use the degree sign (°) to indicate standard reference state conditions and an asterisk (*) for ideality.

To obtain thermodynamic parameters relative to the so defined standard state we normalize the Richards function as

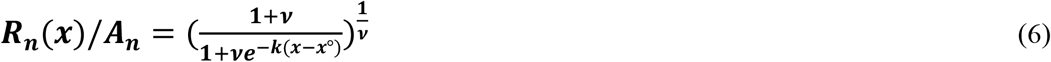

where x is the logarithm of concentration of the titrated molecule and R_n_(x) is the Richards function normalized so that the value of the function is 1 at the point of inflection, where x=x°. In this parametrization ν is the asymmetry parameter in the Richards function and the exponent of the corresponding Richards differential equation (47).

The logistic function defines an ideal reaction where the decrease of free binding partner is symmetric to the increase of the bound form. Microspot immunoassay conditions are non-ideal but rather limiting conditions: Abs bind to the microspot with negligible change in their composition while reaching equilibrium. The Richards function, a form of generalized logistic function, describes non-ideality by allowing asymmetry in these curves and the asymmetry is captured by this additional ν parameter. In physical chemistry the variable that adjusts concentration to thermodynamic activity by accounting for non-ideality in the reaction is the thermodynamic activity coefficient. The activity coefficient therefore changes as the concentration and interactions of Abs change during titration. The observed asymmetric titration curve characterizes the extent of this deviation from ideality.

Overall, in our model the equilibrium concentration of [AbAg] for each [Ab] and [Ag] composition is obtained by the calculation of the standard state concentrations [AbAg]°, [Ab]°, [Ag]° and asymmetry parameters from the estimated values of fitting variables.

The fitting is carried out with the logarithmic forms of the two Richards functions as

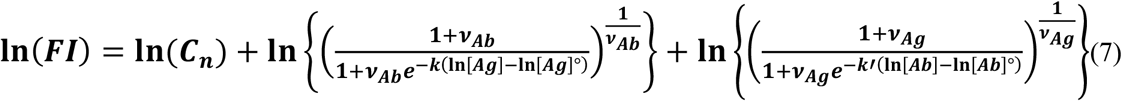

where C_n_ stands for the fluorescence intensity of the complexes in the standard state [AbAg]°. Curve fitting delivers the estimated values for C_n_, ν_Ab_, ν_Ag_, ln[Ag]° and ln[Ab]°. [AbAg]° can be calculated from C_n_ by calibrating the measurement.

### 3.2 Calibration of measurement

Each subarray of the microarray slide contained a dilution series of a mixture of purified human IgM, IgG and IgA, which we used as reference measurement points for calibration. Calibration curves obtained by fitting linear regression curves to log-log datasets (Fig.3A) showed strong and closely linear correlations in this measurement range for IgA (slope=1.05, R^2^=0,99), IgG (slope=0.95, R^2^=0.99) and IgM (slope=0.85, R^2^=0.99). These curves establish the relationship between FI and Ab concentrations and were used for calculating [AbAg]°. Equations for the calculation are available as Supplementary data.

We used the monoclonal Ab AD1.1.10, specific for the hexahistidine tag engineered to the C-terminus of the recombinant RBD, to examine the performance of the measurement system and the fitting strategy with an Ab of known concentration. A series of different concentrations of the Ab was reacted with the microarray and detected using an anti-mouse IgG secondary reagent. The binding data was then fitted using the above algorithm (Fig. 3B-2). The calibration results indicated that the measurement system is suitable to examine Ab reactivity in the low nanomolar to micromolar [Ab] range, using microspotted [Ag] in a similar nanomolar to micromolar range.

**Figure 3.**
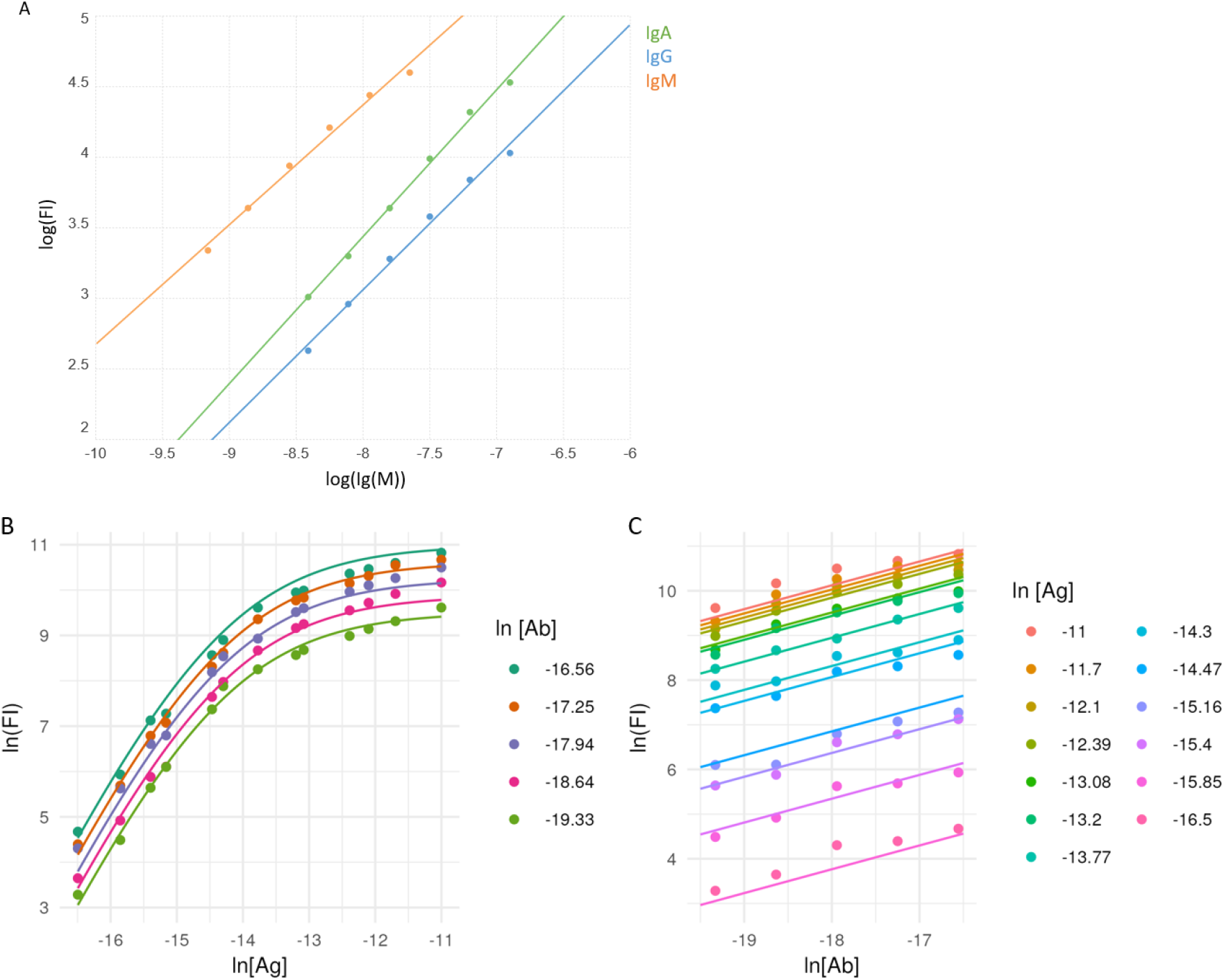
Calibration of Ab signals and curve fitting. A) A mixture of purified immunoglobulins was used to relate fluorescent signals of binding (FI) to molar concentration, using linear regression. Averaged signals of printed Ig mix from all subarrays are represented by dots, lines stand for the fitted function. B) A monoclonal Ab specific for the hexahistidine tag in the recombinant viral RBD was used to characterize the measurement system. Measurement data (dots) and fitted curves for Ag (B) and Ab (C) titration curves are shown.

### 3.3 Complex formation in microspots and in ELISA

When serum Abs are reacted with Ag, the amount of generated AbAg complexes depends on the reaction conditions, primarily the applied concentration of the reactants. This is one of the reasons why different Ab serological tests cannot be compared. By defining the standard reaction conditions as those with standard concentration of both Ab and Ag ([Ab]=[Ab]*, [Ag]=[Ag]°) in a measurement system where Ab is in huge excess, we also define the standard concentration of complexes: [AbAg]°. Its value can be calculated from the calibration curves and ln(C_n_), and represents a reference point from which [AbAg] values deviate depending on ln[Ab] and ln[Ag], as shown in the 3D surface of Figure 1B.

We measured the binding of IgA, IgG and IgM to recombinant SARS-CoV-2 RBD in serum samples from COVID-19 negative and positive individuals. Only samples with sufficient datapoints could be fitted by the algorithm, from the 10 negative samples only those with strong enough binding signals could be fitted (negative group: IgA, n=8; IgG, n=3; IgM, n=3). Two samples from the seropositive set were excluded from the analysis because the delay from COVID-19 onset to blood sampling was too short (<14 days) and accordingly the obtained measurement data were outliers in the group, so the analyzed data contained 18 samples for all three Ab classes which were all fitted successfully (positive group: n=18). There was a significant increase of [AbAg]° values in the positive group that was similar in all three measured Ab classes, reflecting an immune response with increased IgA, IgG and IgM binding to the viral protein in the infected individuals (Fig.4A). We compared the available ELISA results of the positive group to our microarray-derived [AbAg]° data. The logarithm of ELISA units were positively correlated with the logarithm of standard complex concentrations for IgG (R^2^=0.48, p=0.0014) and IgM (R^2^=0.61, p=0.0001) (Fig.4B-C).

**Figure 4.**
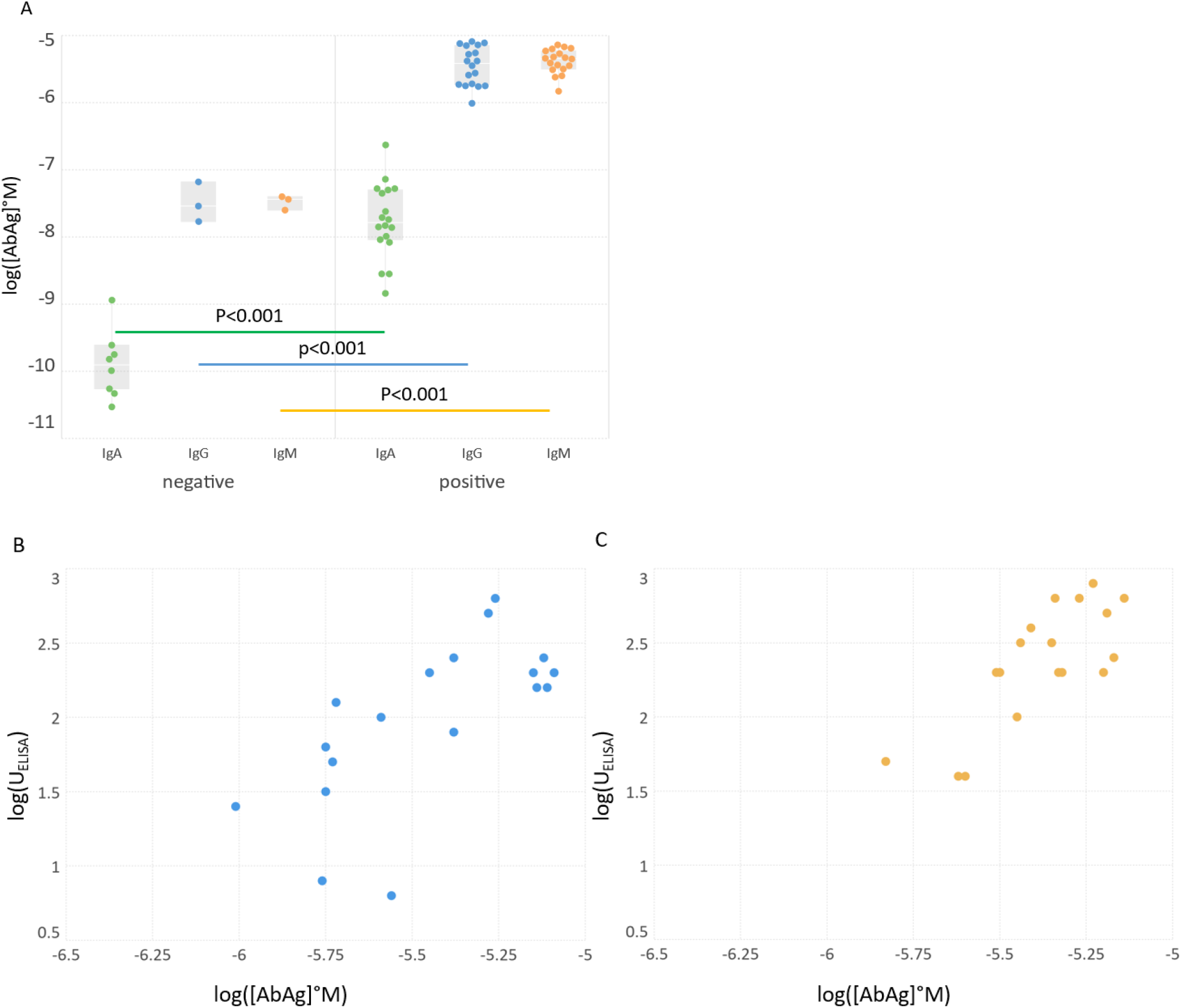
Standard complex concentrations in the groups and their relationship with ELISA results. A) Comparison of standard concentrations in between groups and Ab classes. B-C) Logarithmic values of ELISA results of positive sera were correlated with the obtained log[AbAg]° concentrations. ELISA results for IgA were not available. Significance of differences is characterized by the shown p value above the bars connecting box plots.

### 3.4 Calculation of the affinity and chemical potential of serum Ab

The application of Richards function for data fitting allows for deviation from ideality, which is represented by the logistic function. In our measurement system ideality would be the symmetric increase of bound reactants and decrease of free reactants (Fig.5). The extent of deviation is measured by the asymmetry parameters, which then allow to characterize non-ideality by biochemical variables. The real KD is calculated from the observed apparent KD’_Ag_, defined as being equivalent to the standard Ag concentration [Ag]°. At a distance of In (ν_*Ag*_) from ln[Ag]° is the ideal concentration ln[Ag]* at which half of the binder Ab would be saturated and therefore [Ag]*=KD (Fig.5A). We assume that under equilibrium conditions ideally (in solution) the same concentration of serum Abs would be required to saturate half of the Ag molecules, thus [Ab]*=KD. Under our asymmetric microspot measurement conditions, the point of inflection shifts depending on the effective serum Ab concentration relative to the KD. Thus, when [Ab]=KD then half of Ag is saturated and by diluting the serum we observe a decrease in [AbAg] (Fig.5B). The inflection point is exactly at the undiluted serum measurement point. When [Ab]>KD the inflection point shifts to the left by In (ν_*Ab*_) and falls on the titration curve; when [Ab]<KD we observe no inflection because it falls beyond the titration curve. Thus, asymmetry parameter ν_*Ab*_ is the ratio [Ab]/KD.

**Figure 5.**
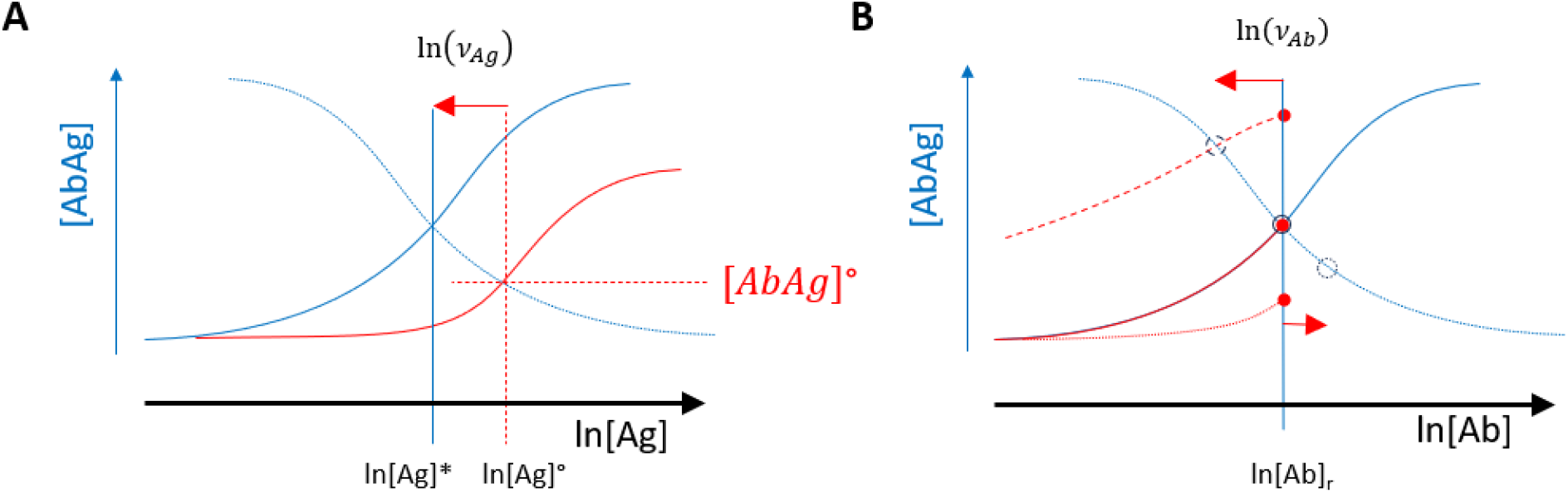
Effects of the asymmetry parameters on affinity and chemical potential. A) The asymmetry parameter of Ag titration defines the distance (red arrow) between the observed inflection point at ln[Ag]° and the inflection point of ideal reaction at ln[Ag]*. Ideal curves of accumulating bound reactant (blue) and decreasing free reactant (slashed line) intersect at ln[Ag]*. B) The asymmetry parameter of Ab titration determines the distance (red arrows) between undiluted serum (filled red circles) and the inflection point of the titration curve (empty blue circles), which is at the intersection with the curve of ideal decrease (slashed line). The value of the asymmetry parameter determines the shape of the curve: when it is bigger than 1, we observe an inflection point (slashed red line), when it is 1 then the curve starts at the point of inflection (solid red line), when it is less than 1, the point of inflection is beyond the measurement range (dotted red line). The value of rate parameter k is assumed 1 and it is therefore omitted from the figure.

The chemical potential of a substance is the slope of the free energy in the system as a function of the concentration of the substance. It is influenced by the composition of the system and the concentration of the substance. If we use ideality as a reference point, chemical potential can be expressed as

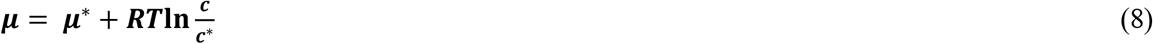

where μ is chemical potential, μ* is chemical potential under ideal conditions, R is universal gas constant, T is thermodynamic temperature, c is molar concentration and c* is concentration under ideal conditions. In our model we can think of the chemical potential of serum Abs as the potential to release free energy by adding Ag to the system. Since Abs binding to any Ag are not a single molecular species this potential reflects an average reactivity. If we calculate the molar free energy change from the KD of the reaction

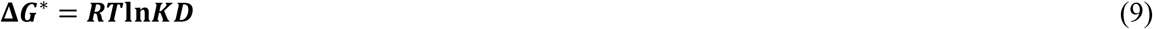

it will characterize the composition of the serum with respect to reactivity against the tested Ag. If this is partial molar free energy change under ideal conditions then, by using the asymmetry parameter of Ab titration we can calculate chemical potential in undiluted serum as

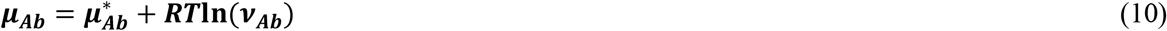

We can now define besides the qualitative parameters of apparent and true KD the quantitative parameter that has universal biochemical units and can characterize serum Ab reactivity. The calculated physico-chemical variables of RBD specific IgA, IgG and IgM in the two groups were compared (Fig.6D). While KD’_Ag_ differences did not reach significance, KD, ν_*Ag*_ and *μ*_*Ab*_ were significantly different for all the Ab classes between the two sample groups.

**Figure 6.**
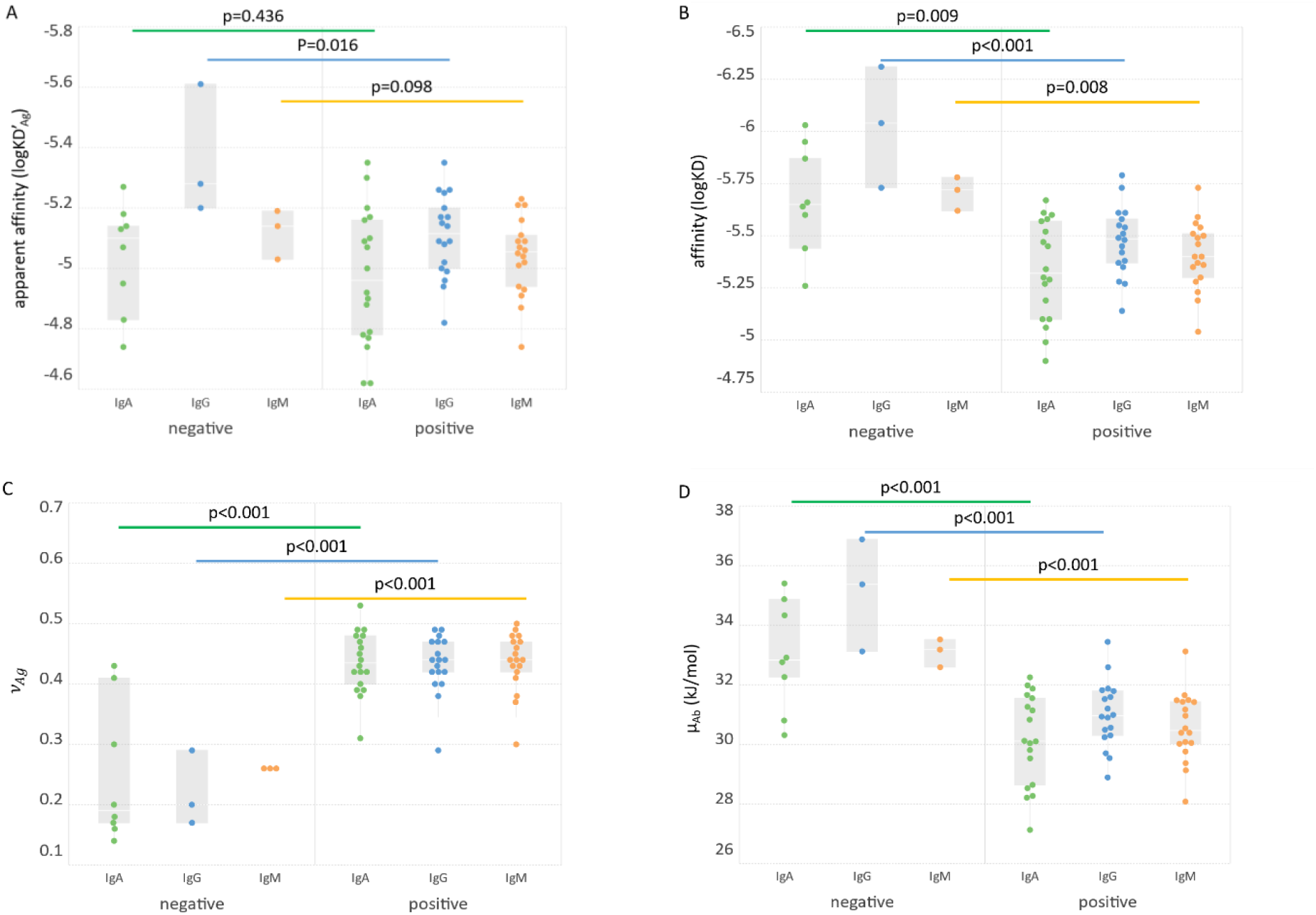
Comparison of groups based on biochemical variables. The apparent affinity (A), affinity (B), asymmetry parameter (C) and chemical potential (D) of three Ab classes in the two groups are shown. Significance of Ab class group differences (bars connecting box plots) is characterized by the shown p values.

## 4. Discussion

Conventional approaches to the qualitative and quantitative characterization of specific serum Abs yield arbitrary units or titers. To obtain biochemically meaningful, truly quantitative results, we applied physical chemistry in the interpretation of microspot Ag titration results. First, we defined the standard chemical thermodynamic state of a particular Ag molecule in a particular serum with respect to a particular Ab class that is measured. The aim of the measurement itself is then the estimation of Ag density in the microspot and the relative Ab concentration of serum Abs required to bring the system to the standard state. If the measurement data are suitable to identify the standard state, we can model the behavior of the system under different conditions.

By employing the combination of two Richards functions, for fitting both Ab and Ag titration curves, we obtain values of the key variables: standard state concentrations and asymmetry parameters. A 3D surface plot of the relative thermodynamic activity of Ab-Ag complexes (Figure 1C) identifies Ab and Ag concentrations at which changes in either of them bring about the greatest change in free energy. On one hand that means the immune system is tuned to sensitively respond to changes in [Ag] in this range. On the other hand, it is this [Ab] that efficiently maintains [Ag] at the set level. Changing [Ag] will trigger B-cell receptor signals in memory B cells, leading to expansion and Ab production (47). Thus, the standard state corresponds to maximal relative thermodynamic activity and represents a steady state the immune system ideally maintains towards this particular Ag. This state is a function of B-cell differentiation, which determines affinity (50), Ag abundance, which drives B-cell proliferation (51), and immune complex removal through Fc receptors, which maintains the flow of Ag towards molecular degradation (52).

A key novelty of our approach is the use of chemical potential for the characterization serum Ab reactivity. In the blood millions of Ab binding sites are present, corresponding to the millions of distinct Ab structures produced in the host organism. Any Ag molecule that comes into contact with serum therefore probes those millions of binding sites; what we observe upon measuring Ab binding is the cumulative result of a distribution of binding strengths. We can no longer speak of distinct concentrations and affinities but rather of the cumulative binding strength and concentration of all binders. Chemical potential is a suitable physico-chemical variable to characterize a heterogenous system since it integrates qualitative 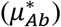 and quantitative (c/c*) aspects of binding into a single variable *μ*_*Ab*_. For a full description of Ab reactivity, the microspotted Ag and the measured Ab isotype should be stated, e.g. *μ*_*IgG*_ (*RBD*_*SARS*−2_).

Serological measurements play a key role during pandemics in several aspects: they allow seroepidemiological monitoring of disease spread, assessment of individual responses against pathogens or vaccines and as a correlate of protection from disease. Multitudes of various assays had been developed recently for SARS-CoV-2 serology, with different platforms, Ags, methods and aims. Neutralization assays are considered good correlates of protection and they show correlations with serological measurements. The determination of Ab affinity has also been suggested to provide useful clinical information (16,53). Unfortunately, most of these assays generate distinct results in terms of units and comparability even after transformations. An assay with truly quantitative readout of chemical properties of SARS-CoV-2 specific Abs could substantially improve our immunological understanding of COVID serology. Our observations on this limited set of serum samples already provide interesting insights. The commercial, conventional technology used here for comparison is the ELISA. The positive correlation between standard complex concentration [AbAg]° and ELISA units (Fig.4B-C) confirms that in spite of different detection methods the generation of bound Ab is similar. ELISA tests are adjusted with respect to coating Ag concentration and secondary reagent so as to characterize serum Ab reactivity in the relevant concentration range, which is around the standard state of our definition. Unexpectedly, the affinity and chemical potential were lower in the COVID-19 positive group in spite of increased [AbAg]° values (Fig.5). This result suggests that the Abs produced during the first weeks of infection are targeted against a wide range of RBD epitopes but bind with poor affinity. This is supported by other studies showing a negative impact of pre-existing common cold coronavirus immunity on SARS-CoV-2 Ab response (54–56). Antigenic sin could therefore be the reason for an increase of bound Ab molecules along with decreased average affinity and chemical potential.

The current limitation of the method with respect to analytical sensitivity is the requirement of a sufficient number of datapoints from the measurement for curve fitting. For weakly reactive (seronegative) serum samples this is a difficult task as it would require a high number of very dense Ag microspots and close to undiluted serum. As long as the characterization of sera that are positive for the given Ag is the goal it may not be a serious shortcoming but it will be safer to work with a method that yields quantitative results even for negative samples. With respect to reproducibility, the accuracy of the Ag density in the microspots is critical. This property is determined and can be controlled during the production of microspot arrays, so it is not influenced by the immunoassay conditions in the laboratory. This study is not a systematic assessment of SARS-CoV-2 specific humoral immune response. The results only show that SARS-CoV-2 specific immune responses can be subjected to a deep thermodynamic analysis using this technology if responses exceed a certain minimal threshold. Here we aimed to demonstrate the concept of application of physical chemistry to quantitative serology via the use of dual-titration and a normalized, generalized logistic function. Further studies using this approach will be needed to reveal the potency of the technology in discovering immunological phenomena not addressed by conventional technologies. We expect that technical improvements and larger scale production of the Ag microspot arrays will increase the efficiency of fitting and render the method suitable for routine use.

Overall, our technology is unique in the sense that it simultaneously estimates all physico-chemical parameters relevant for immunochemical thermodynamic profiling of different Ab classes. The method is based on planar protein microarray technology, which is well-established and available by now in central laboratories. We envisage the application of this technology when a deeper analysis of serological reactivity is needed. In clinical diagnostics it could follow screening steps in the diagnostic algorithm. Once diagnosis is established, the truly quantitative data should be useful for monitoring and adjusting therapy. Beyond medical serology, the generation of Ab binding data with universal units should contribute to the generation of databases for systems immunology in general.

## Supporting information

Supplemental Methods Array protocol

Supplemetal Methods Protein characterization

## Supplementary data

- Mass spectrometry proteomics data of RBD protein is available on PRIDe. LC-MS experiments of SARS-CoV2 Receptor Binding Domain WT. PXD040415.
- Detailed microarray protocol: Detailed characterization of the design, generation and use of microspot immunoassay.
- Biochemical characterization of RBD protein: Description of the biochemical analysis of recombinant SARS-CoV-2 RBD protein quality.
- Data obtained and generated for quantitative serological measurements: files containing all the analyzed data after data cleaning, fitting binding curves and the predicted variables.

## Author contributions

ÁK, TP and JP conceptualized the application of Richards curve to titration data. ÁK, ZH, KP curated the data obtained by measurements. ÁK, KP and DNL carried out formal analysis of immunoassays and protein MS data, respectively. PS and JP arranged funding support. ZH and LB performed experiments. ZH, JZK, DNL and LB contributed to methods development. PS and JP administered the project. JZK, EU and JP provided resources for the project. ÁK worked out the fitting algorithm. EU, TP and JP supervised the project. KP and JP visualized concepts and data. ZH, JZK and JP prepared drafts; KP, TP and JP edited and finalized writing of the manuscript.

## Grant information

No dedicated funding was used for this study.

## Acknowledgments

JZK would like to thank Jonas Heilskov Graversen from the Institute of Molecular Medicine, University of Southern Denmark for the advice within plasmid design and construction.

## Conflict of interest

K.P., Z.H. and J.P. are inventors in a patent application submitted by Diagnosticum Zrt, related to the technology described in this paper.

## Notes

### Competing Interest Statement

K.P., Z.H. and J.P. are inventors in a patent submission by Diagnosticum Zrt, related to the technology described in this paper.

### Summary of Updates

a new set of commercially available serum samples with PCR confirmed disease was analysed and the biochemical interpretation of curve fitting has been improved. As a result the figures show the estimated biochemical units for the new dataset.

## References

1. Cheng HD, Dowell KG, Bailey-Kellogg C, Goods BA, Love JC, Ferrari G, Alter G, Gach J, Forthal DN, Lewis GK, et al. Diverse antiviral IgG effector activities are predicted by unique biophysical antibody features. Retrovirology (2021) 18:35. doi:10.1186/s12977-021-00579-9

2. Chackerian B, Peabody DS. Factors That Govern the Induction of Long-Lived Antibody Responses. Viruses (2020) 12: doi:10.3390/v12010074

3. Choteau M, Scohy A, Messe S, Luyckx M, Dechamps M, Montiel V, Yombi JC, Gruson D, Limaye N, Michiels T, et al. Development of SARS-CoV2 humoral response including neutralizing antibodies is not sufficient to protect patients against fatal infection. Sci Rep (2022) 12:2077. doi:10.1038/s41598-022-06038-5

4. Johnson M, Wagstaffe HR, Gilmour KC, Mai AL, Lewis J, Hunt A, Sirr J, Bengt C, Grandjean L, Goldblatt D. Evaluation of a novel multiplexed assay for determining IgG levels and functional activity to SARS-CoV-2. J Clin Virol (2020) 130:104572. doi:10.1016/j.jcv.2020.104572

5. Farnsworth CW, Case JB, Hock K, Chen RE, O’Halloran JA, Presti R, Goss CW, Rauseo AM, Ellebedy A, Theel ES, et al. Assessment of serological assays for identifying high titer convalescent plasma. medRxiv (2021) doi:10.1101/2021.03.26.21254427

6. Becker M, Strengert M, Junker D, Kaiser PD, Kerrinnes T, Traenkle B, Dinter H, Häring J, Ghozzi S, Zeck A, et al. Exploring beyond clinical routine SARS-CoV-2 serology using MultiCoV-Ab to evaluate endemic coronavirus cross-reactivity. Nat Commun (2021) 12:1152. doi:10.1038/s41467-021-20973-3

7. Klasse PJ. Neutralization of virus infectivity by antibodies: old problems in new perspectives. Advances in Biology (2014) 2014: doi:10.1155/2014/157895

8. Oguntuyo KY, Stevens CS, Hung CT, Ikegame S, Acklin JA, Kowdle SS, Carmichael JC, Chiu H-P, Azarm KD, Haas GD, et al. Quantifying Absolute Neutralization Titers against SARS-CoV-2 by a Standardized Virus Neutralization Assay Allows for Cross-Cohort Comparisons of COVID-19 Sera. MBio (2021) 12: doi:10.1128/mBio.02492-20

9. Khoury DS, Cromer D, Reynaldi A, Schlub TE, Wheatley AK, Juno JA, Subbarao K, Kent SJ, Triccas JA, Davenport MP. Neutralizing antibody levels are highly predictive of immune protection from symptomatic SARS-CoV-2 infection. Nat Med (2021) 27:1205–1211. doi:10.1038/s41591-021-01377-8

10. Prechl J. Why current quantitative serology is not quantitative and how systems immunology could provide solutions. Biologia Futura (2021) doi:10.1007/s42977-020-00061-1

11. Busch NA, Chiew YC, Yarmush ML, Wertheim MS. Development and validation of a simple antigen–antibody model. AIChE J (1995) 41:974–984. doi:10.1002/aic.690410427

12. Berzofsky JA. The assessment of antibody affinity from radioimmunoassay. Clin Chem (1978) 24:419–421.

13. Nisonoff A, Pressman D. Heterogeneity and average combining constants of antibodies from individual rabbits. J Immunol (1958) 80:417–428.

14. Paul WE. Fundamental Immunology. 5th ed. Philadelphia: Lippincott Williams & Wilkins (2003).

15. Lippok S, Seidel SAI, Duhr S, Uhland K, Holthoff H-P, Jenne D, Braun D. Direct detection of antibody concentration and affinity in human serum using microscale thermophoresis. Anal Chem (2012) 84:3523–3530. doi:10.1021/ac202923j

16. Fiedler S, Piziorska MA, Denninger V, Morgunov AS, Ilsley A, Malik AY, Schneider MM, Devenish SRA, Meisl G, Kosmoliaptsis V, et al. Antibody Affinity Governs the Inhibition of SARS-CoV-2 Spike/ACE2 Binding in Patient Serum. ACS Infect Dis (2021) 7:2362–2369. doi:10.1021/acsinfecdis.1c00047

17. Tang C, Verwilligen A, Sadoff J, Brandenburg B, Sneekes-Vriese E, van den Kerkhof T, Dillen L, Rutten L, Juraszek J, Callewaert K, et al. Absolute quantitation of binding antibodies from clinical samples. npj Vaccines (2024) 9:8. doi:10.1038/s41541-023-00793-w

18. Ekins RP. Multi-analyte immunoassay. J Pharm Biomed Anal (1989) 7:155–168. doi:10.1016/0731-7085(89)80079-2

19. Saviranta P, Okon R, Brinker A, Warashina M, Eppinger J, Geierstanger BH. Evaluating sandwich immunoassays in microarray format in terms of the ambient analyte regime. Clin Chem (2004) 50:1907–1920. doi:10.1373/clinchem.2004.037929

20. Papp K, Kovács Á, Orosz A, Hérincs Z, Randek J, Liliom K, Pfeil T, Prechl J. Absolute Quantitation of Serum Antibody Reactivity Using the Richards Growth Model for Antigen Microspot Titration. Sensors (2022)

21. Solastie A, Virta C, Haveri A, Ekström N, Kantele A, Miettinen S, Lempainen J, Jalkanen P, Kakkola L, Dub T, et al. A Highly Sensitive and Specific SARS-CoV-2 Spike- and Nucleoprotein-Based Fluorescent Multiplex Immunoassay (FMIA) to Measure IgG, IgA, and IgM Class Antibodies. Microbiol Spectr (2021) e0113121. doi:10.1128/Spectrum.01131-21

22. Murrell I, Forde D, Zelek W, Tyson L, Chichester L, Palmer N, Jones R, Morgan BP, Moore C. Temporal development and neutralising potential of antibodies against SARS-CoV-2 in hospitalised COVID-19 patients: An observational cohort study. PLoS ONE (2021) 16:e0245382. doi:10.1371/journal.pone.0245382

23. Kubota K, Kitagawa Y, Matsuoka M, Imai K, Orihara Y, Kawamura R, Sakai J, Ishibashi N, Tarumoto N, Takeuchi S, et al. Clinical evaluation of the antibody response in patients with COVID-19 using automated high-throughput immunoassays. Diagn Microbiol Infect Dis (2021) 100:115370. doi:10.1016/j.diagmicrobio.2021.115370

24. Garritsen A, Scholzen A, van den Nieuwenhof DWA, Smits APF, Datema ES, van Galen LS, Kouwijzer MLCE. Two-tiered SARS-CoV-2 seroconversion screening in the Netherlands and stability of nucleocapsid, spike protein domain 1 and neutralizing antibodies. Infect Dis (Lond) (2021) 1–15. doi:10.1080/23744235.2021.1893378

25. Cacaci M, Menchinelli G, Ricci R, De Maio F, Mariotti M, Torelli R, Morandotti GA, Bugli F, Sanguinetti M, Posteraro B. Re-evaluating positive serum samples for SARS-CoV-2 specific IgA and IgG antibodies using an in-house serological assay. Clin Microbiol Infect (2020) doi:10.1016/j.cmi.2020.12.014

26. Sette A, Crotty S. Adaptive immunity to SARS-CoV-2 and COVID-19. Cell (2021) 184:861– 880. doi:10.1016/j.cell.2021.01.007

27. Gaebler C, Wang Z, Lorenzi JCC, Muecksch F, Finkin S, Tokuyama M, Cho A, Jankovic M, Schaefer-Babajew D, Oliveira TY, et al. Evolution of Antibody Immunity to SARS-CoV-2. BioRxiv (2021) doi:10.1101/2020.11.03.367391

28. Qiu M, Shi Y, Guo Z, Chen Z, He R, Chen R, Zhou D, Dai E, Wang X, Si B, et al. Antibody responses to individual proteins of SARS coronavirus and their neutralization activities. Microbes Infect (2005) 7:882–889. doi:10.1016/j.micinf.2005.02.006

29. Heffron AS, McIlwain SJ, Amjadi MF, Baker DA, Khullar S, Sethi AK, Palmenberg AC, Shelef MA, O’Connor DH, Ong IM. The landscape of antibody binding to SARS-CoV-2. BioRxiv (2020) doi:10.1101/2020.10.10.334292

30. Aydillo T, Rombauts A, Stadlbauer D, Aslam S, Abelenda-Alonso G, Escalera A, Amanat F, Jiang K, Krammer F, Carratala J, et al. Immunological imprinting of the antibody response in COVID-19 patients. Nat Commun (2021) 12:3781. doi:10.1038/s41467-021-23977-1

31. Rydyznski Moderbacher C, Ramirez SI, Dan JM, Grifoni A, Hastie KM, Weiskopf D, Belanger S, Abbott RK, Kim C, Choi J, et al. Antigen-Specific Adaptive Immunity to SARS-CoV-2 in Acute COVID-19 and Associations with Age and Disease Severity. Cell (2020) 183:996-1012.e19. doi:10.1016/j.cell.2020.09.038

32. Bölke E, Matuschek C, Fischer JC. Loss of Anti-SARS-CoV-2 Antibodies in Mild Covid-19. N Engl J Med (2020) 383:1694–1695. doi:10.1056/NEJMc2027051

33. Hoepel W, Chen HJ, Geyer CE, Allahverdiyeva S, Manz XD, de Taeye SW, Aman J, Mes L, Steenhuis M, Griffith GR, et al. High titers and low fucosylation of early human anti-SARS-CoV-2 IgG promote inflammation by alveolar macrophages. Sci Transl Med (2021) 13: doi:10.1126/scitranslmed.abf8654

34. Selva KJ, van de Sandt CE, Lemke MM, Lee CY, Shoffner SK, Chua BY, Davis SK, Nguyen THO, Rowntree LC, Hensen L, et al. Systems serology detects functionally distinct coronavirus antibody features in children and elderly. Nat Commun (2021) 12:2037. doi:10.1038/s41467-021-22236-7

35. Atyeo C, Fischinger S, Zohar T, Slein MD, Burke J, Loos C, McCulloch DJ, Newman KL, Wolf C, Yu J, et al. Distinct Early Serological Signatures Track with SARS-CoV-2 Survival. Immunity (2020) 53:524-532.e4. doi:10.1016/j.immuni.2020.07.020

36. Dan JM, Mateus J, Kato Y, Hastie KM, Yu ED, Faliti CE, Grifoni A, Ramirez SI, Haupt S, Frazier A, et al. Immunological memory to SARS-CoV-2 assessed for up to 8 months after infection. Science (2021) 371: doi:10.1126/science.abf4063

37. Wajnberg A, Amanat F, Firpo A, Altman DR, Bailey MJ, Mansour M, McMahon M, Meade P, Mendu DR, Muellers K, et al. Robust neutralizing antibodies to SARS-CoV-2 infection persist for months. Science (2020) 370:1227–1230. doi:10.1126/science.abd7728

38. Nam M, Seo JD, Moon H-W, Kim H, Hur M, Yun Y-M. Evaluation of Humoral Immune Response after SARS-CoV-2 Vaccination Using Two Binding Antibody Assays and a Neutralizing Antibody Assay. Microbiol Spectr (2021) 9:e0120221. doi:10.1128/Spectrum.01202-21

39. Struck F, Schreiner P, Staschik E, Wochinz-Richter K, Schulz S, Soutschek E, Motz M, Bauer G. Vaccination versus infection with SARS-CoV-2: Establishment of a high avidity IgG response versus incomplete avidity maturation. J Med Virol (2021) 93:6765–6777. doi:10.1002/jmv.27270

40. Chen Y, Yin S, Tong X, Tao Y, Ni J, Pan J, Li M, Wan Y, Mao M, Xiong Y, et al. Dynamic SARS-CoV-2-specific B-cell and T-cell responses following immunization with an inactivated COVID-19 vaccine. Clin Microbiol Infect (2022) 28:410–418. doi:10.1016/j.cmi.2021.10.006

41. Wu K, Werner AP, Koch M, Choi A, Narayanan E, Stewart-Jones GBE, Colpitts T, Bennett H, Boyoglu-Barnum S, Shi W, et al. Serum Neutralizing Activity Elicited by mRNA-1273 Vaccine - Preliminary Report. N Engl J Med (2021) doi:10.1056/NEJMc2102179

42. Pegu A, O’Connell S, Schmidt SD, O’Dell S, Talana CA, Lai L, Albert J, Anderson E, Bennett H, Corbett KS, et al. Durability of mRNA-1273 vaccine–induced antibodies against SARS-CoV-2 variants. Science (2021)

43. Ebanks D, Faustini S, Shields A, Parry H, Moss P, Plant T, Richter A, Drayson M. Cross reactivity of serological response to SARS-CoV-2 vaccination with viral variants of concern detected by lateral flow immunoassays. J Infect (2021) doi:10.1016/j.jinf.2021.07.020

44. Kritikos A, Gabellon S, Pagani J-L, Monti M, Bochud P-Y, Manuel O, Coste A, Greub G, Perreau M, Pantaleo G, et al. Anti-SARS-CoV-2 Titers Predict the Severity of COVID-19. Viruses (2022) 14: doi:10.3390/v14051089

45. Muecksch F, Wise H, Templeton K, Batchelor B, Squires M, McCance K, Jarvis L, Malloy K, Furrie E, Richardson C, et al. Longitudinal variation in SARS-CoV-2 antibody levels and emergence of viral variants: implications for the ability of serological assays to predict immunity. medRxiv (2021) doi:10.1101/2021.07.02.21259939

46. Bauer G. The potential significance of high avidity immunoglobulin G (IgG) for protective immunity towards SARS-CoV-2. Int J Infect Dis (2021) 106:61–64. doi:10.1016/j.ijid.2021.01.061

47. Prechl J, Papp K, Kovács Á, Pfeil T. The binding landscape of serum antibodies: how physical and mathematical concepts can advance systems immunology. Antibodies (Basel) (2022) 11:43. doi:10.3390/antib11030043

48. Rappsilber J, Mann M, Ishihama Y. Protocol for micro-purification, enrichment, prefractionation and storage of peptides for proteomics using StageTips. Nat Protoc (2007) 2:1896–1906. doi:10.1038/nprot.2007.261

49. Pfeil T, Herbály B. A linear model for polyclonal antibody–antigen reactions. Math Comput Simul (2022) 198:20–30. doi:10.1016/j.matcom.2022.02.004

50. Prechl J. A generalized quantitative antibody homeostasis model: regulation of B-cell development by BCR saturation and novel insights into bone marrow function. Clin Transl Immunology (2017) 6:e130. doi:10.1038/cti.2016.89

51. Prechl J. A generalized quantitative antibody homeostasis model: antigen saturation, natural antibodies and a quantitative antibody network. Clin Transl Immunology (2017) 6:e131. doi:10.1038/cti.2016.90

52. Prechl J. A generalized quantitative antibody homeostasis model: maintenance of global antibody equilibrium by effector functions. Clin Transl Immunology (2017) 6:e161. doi:10.1038/cti.2017.50

53. Bauer G, Struck F, Staschik E, Maile J, Wochinz-Richter K, Motz M, Soutschek E. Differential avidity determination of IgG directed towards the receptor-binding domain (RBD) of SARS-CoV-2 wild-type and its variants in one assay: Rational tool for the assessment of protective immunity. J Med Virol (2022) doi:10.1002/jmv.28006

54. Lin C-Y, Wolf J, Brice DC, Sun Y, Locke M, Cherry S, Castellaw AH, Wehenkel M, Crawford JC, Zarnitsyna VI, et al. Pre-existing humoral immunity to human common cold coronaviruses negatively impacts the protective SARS-CoV-2 antibody response. Cell Host Microbe (2022) 30:83-96.e4. doi:10.1016/j.chom.2021.12.005

55. Anderson EM, Li SH, Awofolaju M, Eilola T, Goodwin E, Bolton MJ, Gouma S, Manzoni TB, Hicks P, Goel RR, et al. SARS-CoV-2 infections elicit higher levels of original antigenic sin antibodies compared with SARS-CoV-2 mRNA vaccinations. Cell Rep (2022) 41:111496. doi:10.1016/j.celrep.2022.111496

56. Aguilar-Bretones M, Westerhuis BM, Raadsen MP, de Bruin E, Chandler FD, Okba NM, Haagmans BL, Langerak T, Endeman H, van den Akker JP, et al. Seasonal coronavirus-specific B cells with limited SARS-CoV-2 cross-reactivity dominate the IgG response in severe COVID-19. J Clin Invest (2021) 131: doi:10.1172/JCI150613

