## Supplemental Methods Array protocol for "Deep physico-chemical characterization of the serum antibody response using a dual-titration microspot assay"

Supplementary data: Deep physico-chemical characterization of individual serum antibody responses against SARS-CoV-2 RBD using a dual titration microspot assay ÁK, ZH, KP, JZK, DNL, PS, LB, EU, TP, JP

### Production and use of antigen microarrays

#### Microarray printing protocol

Printing was performed with a BioRad Calligrapher MiniArrayer, using 2 solid pins. Materials were printed on NHS- coated Nexterion H glass slides from Schott-Nexterion. Antigens and calibration immunoglobulin mix were prepared and diluted in printing buffer, containing 0,01% Tween20 + 2% glycerol + 0,5% DMSO in phosphate-buffered saline.

A 7x7 matrix was printed in each subarray in a 16-pad format. A 6x7 microspot matrix comprised 14 different dilutions of the antigen, each with 3 parallel spots. A 1x7 microspot row was used for the immunoglobulin reference for calibration.

Antigens printed:

1. **SARS-CoV2 Spike RBD 319-541 (OvodanBiotech)**, „RBDwuh"

The 14-point dilution series was made with a combination of a ½ and ⅓ diluting series. The final concentrations were as follows (μM)

| {1} | {2} | {3} | {4} | {5} | {6} | {7} | {8} | {9} | {10} | {11} | {12} | {13} | {14} |
|---|---|---|---|---|---|---|---|---|---|---|---|---|---|
| 16.666 | 8.333 | 5.555 | 4.166 | 2.083 | 1.851 | 1.041 | 0.617 | 0.520 | 0.260 | 0.205 | 0.130 | 0.068 | 0.065 |

Antibodies printed:

2. **IgA-IgM-IgG antibody mix ('AMG') for calibration:**
   - **human IgA** (Jackson, 009-000-011)
   - **human IgM** (Sigma, I8260)
   - **human IgG** (Sigma, I2511)

The mix contains a final concentration of 10 µg/ml for each antibody. This was the 1st point of a 7-point ½-diluting series.

Slide layout with subarrays numbered from 1 to 16:

| | | | | | | | |
|---|---|---|---|---|---|---|---|
| 2 | 4 | 6 | 8 | 10 | 12 | 14 | 16 |
| 1 | 3 | 5 | 7 | 9 | 11 | 13 | 15 |

*

Subarray layout within slide

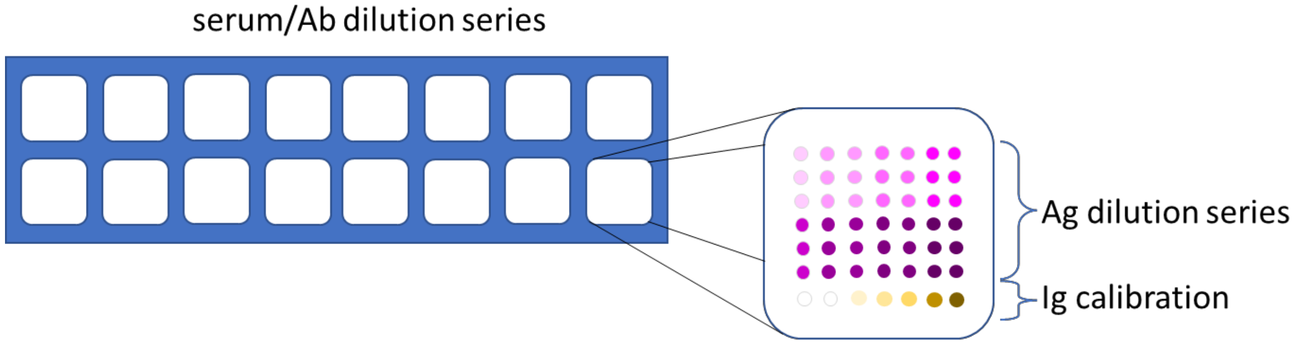


„RBD wuhan” subarray arrangement (1 subarray enlarged):

| | | | | | | | |
|---|---|---|---|---|---|---|---|
| RBDwuh{14} | RBDwuh{13} | RBDwuh{12} | RBDwuh{11} | | RBDwuh{10} | RBDwuh{9} | RBDwuh{8} |
| RBDwuh{14} | RBDwuh{13} | RBDwuh{12} | RBDwuh{11} | | RBDwuh{10} | RBDwuh{9} | RBDwuh{8} |
| RBDwuh{14} | RBDwuh{13} | RBDwuh{12} | RBDwuh{11} | | RBDwuh{10} | RBDwuh{9} | RBDwuh{8} |
| RBDwuh{7} | RBDwuh{6} | RBDwuh{5} | RBDwuh{4} | | RBDwuh{3} | RBDwuh{2} | RBDwuh{1} |
| RBDwuh{7} | RBDwuh{6} | RBDwuh{5} | RBDwuh{4} | | RBDwuh{3} | RBDwuh{2} | RBDwuh{1} |
| RBDwuh{7} | RBDwuh{6} | RBDwuh{5} | RBDwuh{4} | | RBDwuh{3} | RBDwuh{2} | RBDwuh{1} |
| AMG{7} | AMG{6} | AMG{5} | AMG{4} | | AMG{3} | AMG{2} | AMG{1} |

*

Following printing slides were dried at 37°C for 60 minutes then blocked at 37°C for 60 minutes in Tris 0.1M (pH=8.0) buffer. When finished, slides were quickly rinsed in distilled water twice, then dried for 2 minutes in a fix-angled slide centrifuge. Finally, they were sealed in a light-protected bag and kept on 4°C until usage.

#### Microspot immunoassay protocol

Slides were taken out from 4°C and allowed to reach room temperature on the bench for 30 minutes. Slides were fitted in a Grace Biolabs steel clipped 16-chamber system for the immunoassay. Slides were rehydrated in PBS for 3x10 minutes, then blocked in PBS containing 2% bovine serum albumin for 30 minutes at 37°C. Reaction chambers were then incubated with sera of different dilutions (sample buffer 0.5% bovine serum albumin, 0.05% Tween20 in PBS), the two slides were incubated with the sera in a completely identical way.

A typical slide was assembled as follows (numbers stand for dilution factor):

| | | | | | | | |
|---|---|---|---|---|---|---|---|
| spl_buffer | spl_buffer | ser_5x | ser_5x | ser_25x | ser_25x | ser_125x | ser_125x |
| ser_625x | ser_625x | ser_10x | ser_10x | ser_100x | ser_100x | ser_1000x | ser_1000x |

Slides were washed 3x5 minutes in PBS-0.05%Tween20 on an orbital shaker.

The detection of the different Ig classes was performed with the following secondary labeled antibody mixture:

a. anti-human-**IgA** – A647 (Jackson, #:109606011, L:152435, dil:2021.02.10.) 1/1000
b. anti-human-**IgM** – Cy3 (Jackson, #:109166129, L:90910) 1/2000
c. anti-human **IgG** F(ab')$_2$ – A488 (Jackson, #:109646097, L:105228) 1/1000

Reaction layout for heavy chain detection

| | | | | | | | |
|---|---|---|---|---|---|---|---|
| AMG_mix | AMG_mix | AMG_mix | AMG_mix | AMG_mix | AMG_mix | AMG_mix | AMG_mix |
| AMG_mix | AMG_mix | AMG_mix | AMG_mix | AMG_mix | AMG_mix | AMG_mix | AMG_mix |

Incubations with the labeled antibodies diluted in sample buffer were at RT for 30 minutes.

Slides were then washed 3 x 10 min in PBS-0.05% Tween20, then completely dryed before being scanned with a Sensovation FLAIR microarray reader for the ’Blue’ channel (488nm) and a Sensovation SensoSpot microarray reader for the ’Green’ (532nm) and the ’Red’ channels (647nm).
