## Supplementary material for "Deep physico-chemical characterization of the serum antibody response using a dual-titration microspot assay": Supplemetal Methods Protein characterization

Extended data: Deep physico-chemical characterization of individual serum antibody responses against SARS-CoV-2 RBD using a dual titration microspot assay ÁK, ZH, KP, JZK, DNL, PS, LB, EU, TP, JP

**SARS-CoV-2 Spike RBD 319-541 protein characterization.**

Sodium Dodecyl Sulfate Polyacrylamide gel electrophoresis indicated a major protein band at 37kDa (Figure XXX A) whereas the theoretical molecular weight was 27 kDa. The 9 kDa difference was associated with the two glycosylation sites on NIT (42N) and NAT (54N) (Figure XXX B) indicated by the mass spectrometric analysis. Full sequence coverage was achieved, by searching the mass spectrometric data against a peptide library of the major variants of Sars-CoV-2 Spike RBD 319-541 with a His-Tag at the C-terminal. The searches were performed on both non-de-glycosylated and de-glycosylated samples. Nearly full sequence coverage was achieved with the non-de-glycosylated sample, the peptide missing was FP**NIT**NLCPFGEVF**NAT**, where two potential glycosylation is located, marked in bold. Coverage across this sequence was achieved by de-glycosylation of the sample, shown in Figure XXXB. Furthermore, deamination of both N (IT) and N (AT) are observed for the de-glycosylated sample, thereby proving that glycosylation is present at both sites in the sample.

Supplemetary Figure RBD .

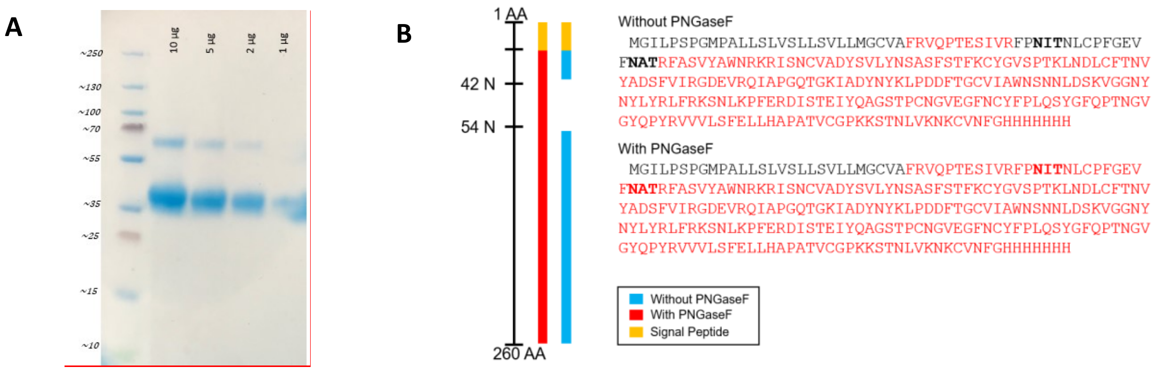
