## Supplementary material for "Deep physico-chemical characterization of the serum antibody response using a dual-titration microspot assay": Supplemetal data

| ID | Group | Antibody | Parameter | Value | 2.5% | 97.5% |
|---|---|---|---|---|---|---|
| PS303 | positive | IgA | $\ln(C_n)$ | 9,674618 | 8,947996 | 10,51363 |
| PS303 | positive | IgA | ν | 0,488064 | 0,46123 | 0,512299 |
| PS303 | positive | IgA | ln[Ag]° | -10,9237 | -11,3364 | -10,4569 |
| PS305 | positive | IgA | $\ln(C_n)$ | 9,960744 | 9,15654 | 10,89315 |
| PS305 | positive | IgA | ν | 0,483721 | 0,452661 | 0,511354 |
| PS305 | positive | IgA | ln[Ag]° | -11,0066 | -11,4414 | -10,5048 |
| PS308 | positive | IgA | $\ln(C_n)$ | 10,01634 | 9,545236 | 10,51714 |
| PS308 | positive | IgA | ν | 0,43065 | 0,408351 | 0,451348 |
| PS308 | positive | IgA | ln[Ag]° | -11,2904 | -11,5371 | -11,0323 |
| PS310 | positive | IgA | $\ln(C_n)$ | 8,456381 | 7,948173 | 8,987007 |
| PS310 | positive | IgA | ν | 0,420634 | 0,392579 | 0,446183 |
| PS310 | positive | IgA | ln[Ag]° | -11,9647 | -12,2683 | -11,6643 |
| PS326 | positive | IgA | $\ln(C_n)$ | 7,782106 | 6,019582 | 10,38916 |
| PS326 | positive | IgA | ν | 0,42498 | 0,326973 | 0,492988 |
| PS326 | positive | IgA | ln[Ag]° | -11,2579 | -12,5325 | -9,73128 |
| PS329 | positive | IgA | $\ln(C_n)$ | 11,23564 | 10,70028 | 11,80037 |
| PS329 | positive | IgA | ν | 0,30997 | 0,269921 | 0,346368 |
| PS329 | positive | IgA | ln[Ag]° | -11,7144 | -11,9359 | -11,4782 |
| PS330 | positive | IgA | $\ln(C_n)$ | 10,28675 | 9,512029 | 11,14819 |
| PS330 | positive | IgA | ν | 0,391409 | 0,34822 | 0,429252 |
| PS330 | positive | IgA | ln[Ag]° | -11,2396 | -11,6178 | -10,824 |
| PS331 | positive | IgA | $\ln(C_n)$ | 12,65611 | 12,18983 | 13,13517 |
| PS331 | positive | IgA | ν | 0,479216 | 0,454343 | 0,501903 |
| PS331 | positive | IgA | ln[Ag]° | -12,3278 | -12,5812 | -12,0739 |
| PS332 | positive | IgA | $\ln(C_n)$ | 11,19917 | 10,75622 | 11,6641 |
| PS332 | positive | IgA | ν | 0,456948 | 0,437507 | 0,475121 |
| PS332 | positive | IgA | ln[Ag]° | -11,517 | -11,7436 | -11,2798 |
| PS352 | positive | IgA | $\ln(C_n)$ | | | |
| PS352 | positive | IgA | ν | | | |
| PS352 | positive | IgA | ln[Ag]° | | | |
| PS353 | positive | IgA | $\ln(C_n)$ | 11,54348 | 10,6637 | 12,62624 |
| PS353 | positive | IgA | ν | 0,527471 | 0,499826 | 0,55221 |
| PS353 | positive | IgA | ln[Ag]° | -10,6378 | -11,1175 | -10,0382 |
| PS601 | positive | IgA | $\ln(C_n)$ | 10,30619 | 10,08686 | 10,52893 |
| PS601 | positive | IgA | ν | 0,494851 | 0,485849 | 0,503536 |
| PS601 | positive | IgA | ln[Ag]° | -12,1742 | -12,2999 | -12,0481 |
| PS604 | positive | IgA | $\ln(C_n)$ | 8,445754 | 7,648064 | 9,342494 |
| PS604 | positive | IgA | ν | 0,416711 | 0,374279 | 0,453542 |
| PS604 | positive | IgA | ln[Ag]° | -11,3194 | -11,7688 | -10,837 |
| PS607 | positive | IgA | $\ln(C_n)$ | 11,07 | 10,17792 | 12,03643 |
| PS607 | positive | IgA | ν | 0,398731 | 0,341399 | 0,446923 |
| PS607 | positive | IgA | ln[Ag]° | -11,674 | -12,0921 | -11,2235 |
| PS609 | positive | IgA | $\ln(C_n)$ | 7,870312 | 7,355029 | 8,424594 |
| PS609 | positive | IgA | ν | 0,361861 | 0,330124 | 0,390655 |
| PS609 | positive | IgA | ln[Ag]° | -11,2672 | -11,5345 | -10,9876 |
| PS610 | positive | IgA | $\ln(C_n)$ | 9,973343 | 9,370651 | 10,61467 |

| | | | | | | |
|---|---|---|---|---|---|---|
| PS610 | positive | IgA | ν | 0,445595 | 0,416749 | 0,471732 |
| PS610 | positive | IgA | ln[Ag]° | -11,7438 | -12,0839 | -11,3933 |
| PS611 | positive | IgA | $\ln(C_n)$ | 9,46926 | 8,726852 | 10,34182 |
| PS611 | positive | IgA | ν | 0,421287 | 0,390936 | 0,448695 |
| PS611 | positive | IgA | ln[Ag]° | -10,6297 | -11,0132 | -10,1862 |
| PS619 | positive | IgA | $\ln(C_n)$ | 11,23364 | 10,67503 | 11,81577 |
| PS619 | positive | IgA | ν | 0,383404 | 0,346401 | 0,416486 |
| PS619 | positive | IgA | ln[Ag]° | -11,8922 | -12,1514 | -11,6248 |
| PS620 | positive | IgA | $\ln(C_n)$ | 10,47893 | 10,11737 | 10,8511 |
| PS620 | positive | IgA | ν | 0,441887 | 0,423942 | 0,458758 |
| PS620 | positive | IgA | ln[Ag]° | -11,897 | -12,0778 | -11,7118 |
| PS623 | positive | IgA | $\ln(C_n)$ | 7,710879 | 6,948681 | 8,574926 |
| PS623 | positive | IgA | ν | 0,38727 | 0,348711 | 0,421274 |
| PS623 | positive | IgA | ln[Ag]° | -11,0657 | -11,507 | -10,5995 |
| PS326R | positive | IgA | $\ln(C_n)$ | 9,563422 | 9,007638 | 10,16891 |
| PS326R | positive | IgA | ν | 0,470329 | 0,444408 | 0,493907 |
| PS326R | positive | IgA | ln[Ag]° | -10,9931 | -11,2894 | -10,6723 |
| PS352R | positive | IgA | $\ln(C_n)$ | 4,906816 | 4,59337 | 5,243248 |
| PS352R | positive | IgA | ν | 0,12296 | 0,069855 | 0,170361 |
| PS352R | positive | IgA | ln[Ag]° | -11,5126 | -11,6957 | -11,337 |
| PN11 | negative | IgA | $\ln(C_n)$ | 5,810595 | 5,094723 | 6,64403 |
| PN11 | negative | IgA | ν | 0,301774 | 0,248447 | 0,347952 |
| PN11 | negative | IgA | ln[Ag]° | -10,9224 | -11,4096 | -10,4461 |
| PN12 | negative | IgA | $\ln(C_n)$ | 6,117705 | 5,048989 | 7,429271 |
| PN12 | negative | IgA | ν | 0,407877 | 0,32264 | 0,473844 |
| PN12 | negative | IgA | ln[Ag]° | -12,1456 | -13,0288 | -11,1961 |
| PN13 | negative | IgA | $\ln(C_n)$ | | | |
| PN13 | negative | IgA | ν | | | |
| PN13 | negative | IgA | ln[Ag]° | | | |
| PN14 | negative | IgA | $\ln(C_n)$ | | | |
| PN14 | negative | IgA | ν | | | |
| PN14 | negative | IgA | ln[Ag]° | | | |
| PN15 | negative | IgA | $\ln(C_n)$ | 5,151891 | 4,871554 | 5,452665 |
| PN15 | negative | IgA | ν | 0,114379 | 0,063476 | 0,16022 |
| PN15 | negative | IgA | ln[Ag]° | -11,926 | -12,1237 | -11,7419 |
| PN16 | negative | IgA | $\ln(C_n)$ | 4,547801 | 3,857676 | 5,372452 |
| PN16 | negative | IgA | ν | 0,168239 | 0,073398 | 0,245486 |
| PN16 | negative | IgA | ln[Ag]° | -11,1147 | -11,6263 | -10,6409 |
| PN17 | negative | IgA | $\ln(C_n)$ | 7,601619 | 7,192809 | 8,034438 |
| PN17 | negative | IgA | ν | 0,431074 | 0,408184 | 0,452219 |
| PN17 | negative | IgA | ln[Ag]° | -11,6762 | -11,9419 | -11,4084 |
| PN18 | negative | IgA | $\ln(C_n)$ | 4,097957 | 3,60959 | 4,62301 |
| PN18 | negative | IgA | ν | 0,156009 | 0,072725 | 0,225547 |
| PN18 | negative | IgA | ln[Ag]° | -11,8427 | -12,5951 | -11,3866 |
| PN19 | negative | IgA | $\ln(C_n)$ | 5,565713 | 5,298625 | 5,844204 |
| PN19 | negative | IgA | ν | 0,167991 | 0,127323 | 0,205249 |
| PN19 | negative | IgA | ln[Ag]° | -11,8242 | -12,0274 | -11,6408 |

| | | | | | | |
|---|---|---|---|---|---|---|
| PN20 | negative | IgA | $\ln(C_n)$ | 4,695454 | 3,987723 | 5,49647 |
| PN20 | negative | IgA | $\nu$ | 0,201051 | 0,115264 | 0,271685 |
| PN20 | negative | IgA | ln[Ag]° | -11,3912 | -12,0112 | -10,8966 |

$\ln(C_n)$

| ID | Group | Antibody | Parameter | Value | 2.5% | 97.5% |
|---|---|---|---|---|---|---|
| PS303 | positive | IgG | $\ln(C_n)$ | 10,41268 | 9,551573 | 11,41273 |
| PS303 | positive | IgG | ν | 0,469793 | 0,435144 | 0,50043 |
| PS303 | positive | IgG | ln[Ag]° | -11,0907 | -11,58 | -10,5402 |
| PS305 | positive | IgG | $\ln(C_n)$ | 11,94416 | 11,50372 | 12,40556 |
| PS305 | positive | IgG | ν | 0,45247 | 0,432344 | 0,471251 |
| PS305 | positive | IgG | ln[Ag]° | -11,567 | -11,7951 | -11,3296 |
| PS308 | positive | IgG | $\ln(C_n)$ | 12,01917 | 11,32051 | 12,76977 |
| PS308 | positive | IgG | ν | 0,416857 | 0,378723 | 0,450702 |
| PS308 | positive | IgG | ln[Ag]° | -11,5237 | -11,8376 | -11,1801 |
| PS310 | positive | IgG | $\ln(C_n)$ | 11,32124 | 10,56076 | 12,13329 |
| PS310 | positive | IgG | ν | 0,400405 | 0,353584 | 0,441008 |
| PS310 | positive | IgG | ln[Ag]° | -11,7029 | -12,0352 | -11,3411 |
| PS326 | positive | IgG | $\ln(C_n)$ | 9,223363 | 8,540108 | 9,955402 |
| PS326 | positive | IgG | ν | 0,429061 | 0,391676 | 0,462171 |
| PS326 | positive | IgG | ln[Ag]° | -11,7041 | -12,0827 | -11,3153 |
| PS329 | positive | IgG | $\ln(C_n)$ | 11,61748 | 11,13008 | 12,12242 |
| PS329 | positive | IgG | ν | 0,293425 | 0,250628 | 0,331847 |
| PS329 | positive | IgG | ln[Ag]° | -12,0992 | -12,2934 | -11,896 |
| PS330 | positive | IgG | $\ln(C_n)$ | 11,89227 | 11,42243 | 12,37805 |
| PS330 | positive | IgG | ν | 0,421268 | 0,394414 | 0,445855 |
| PS330 | positive | IgG | ln[Ag]° | -11,9131 | -12,1275 | -11,6908 |
| PS331 | positive | IgG | $\ln(C_n)$ | 11,55506 | 11,17063 | 11,94795 |
| PS331 | positive | IgG | ν | 0,421299 | 0,396179 | 0,44444 |
| PS331 | positive | IgG | ln[Ag]° | -12,3308 | -12,5197 | -12,1404 |
| PS332 | positive | IgG | $\ln(C_n)$ | 11,97363 | 11,65337 | 12,30027 |
| PS332 | positive | IgG | ν | 0,44243 | 0,425432 | 0,458454 |
| PS332 | positive | IgG | ln[Ag]° | -12,1057 | -12,262 | -11,9475 |
| PS352 | positive | IgG | $\ln(C_n)$ | | | |
| PS352 | positive | IgG | ν | | | |
| PS352 | positive | IgG | ln[Ag]° | | | |
| PS353 | positive | IgG | $\ln(C_n)$ | 10,86795 | 10,12122 | 11,68186 |
| PS353 | positive | IgG | ν | 0,441226 | 0,40507 | 0,47331 |
| PS353 | positive | IgG | ln[Ag]° | -11,4986 | -11,8745 | -11,0896 |
| PS601 | positive | IgG | $\ln(C_n)$ | 9,776456 | 9,400764 | 10,16458 |
| PS601 | positive | IgG | ν | 0,477482 | 0,460713 | 0,49323 |
| PS601 | positive | IgG | ln[Ag]° | -11,9111 | -12,1427 | -11,6808 |
| PS604 | positive | IgG | $\ln(C_n)$ | 11,30948 | 10,38562 | 12,36091 |
| PS604 | positive | IgG | ν | 0,490218 | 0,454111 | 0,521818 |
| PS604 | positive | IgG | ln[Ag]° | -11,4238 | -11,918 | -10,8622 |
| PS607 | positive | IgG | $\ln(C_n)$ | 10,49179 | 9,861106 | 11,15408 |
| PS607 | positive | IgG | ν | 0,395659 | 0,355889 | 0,430922 |
| PS607 | positive | IgG | ln[Ag]° | -11,8336 | -12,1167 | -11,5351 |
| PS609 | positive | IgG | $\ln(C_n)$ | 5,428819 | 4,661805 | 6,230433 |
| PS609 | positive | IgG | ν | 0,251911 | 0,149881 | 0,332385 |
| PS609 | positive | IgG | ln[Ag]° | -13,2999 | -14,2417 | -12,6903 |
| PS610 | positive | IgG | $\ln(C_n)$ | 10,46861 | 9,751602 | 11,23372 |

| | | | | | | |
|---|---|---|---|---|---|---|
| PS610 | positive | IgG | ν | 0,431932 | 0,39273 | 0,466408 |
| PS610 | positive | IgG | ln[Ag]° | -11,7159 | -12,0641 | -11,3449 |
| PS611 | positive | IgG | $\ln(C_n)$ | 10,40061 | 9,948266 | 10,88056 |
| PS611 | positive | IgG | ν | 0,466079 | 0,447297 | 0,483591 |
| PS611 | positive | IgG | ln[Ag]° | -11,3875 | -11,6413 | -11,1227 |
| PS619 | positive | IgG | $\ln(C_n)$ | 11,18196 | 10,62573 | 11,76284 |
| PS619 | positive | IgG | ν | 0,382594 | 0,346897 | 0,41469 |
| PS619 | positive | IgG | ln[Ag]° | -11,8932 | -12,1454 | -11,6298 |
| PS620 | positive | IgG | $\ln(C_n)$ | 11,90467 | 11,5228 | 12,29574 |
| PS620 | positive | IgG | ν | 0,440302 | 0,419761 | 0,459447 |
| PS620 | positive | IgG | ln[Ag]° | -12,1072 | -12,2938 | -11,9177 |
| PS623 | positive | IgG | $\ln(C_n)$ | 10,80444 | 10,26219 | 11,37605 |
| PS623 | positive | IgG | ν | 0,470621 | 0,446629 | 0,492608 |
| PS623 | positive | IgG | ln[Ag]° | -11,7152 | -11,9985 | -11,4181 |
| PS326R | positive | IgG | $\ln(C_n)$ | 10,45879 | 10,11005 | 10,81597 |
| PS326R | positive | IgG | ν | 0,491146 | 0,47266 | 0,508379 |
| PS326R | positive | IgG | ln[Ag]° | -11,9575 | -12,1624 | -11,7538 |
| PS352R | positive | IgG | $\ln(C_n)$ | 5,793383 | 5,280809 | 6,334862 |
| PS352R | positive | IgG | ν | 0,275537 | 0,213866 | 0,328167 |
| PS352R | positive | IgG | ln[Ag]° | -12,4046 | -12,7279 | -12,0962 |
| PN11 | negative | IgG | $\ln(C_n)$ | 6,035505 | 5,682441 | 6,408528 |
| PN11 | negative | IgG | ν | 0,173924 | 0,123148 | 0,219328 |
| PN11 | negative | IgG | ln[Ag]° | -12,1661 | -12,3785 | -11,9599 |
| PN12 | negative | IgG | $\ln(C_n)$ | 5,454151 | 4,708471 | 6,276499 |
| PN12 | negative | IgG | ν | 0,203361 | 0,085926 | 0,29488 |
| PN12 | negative | IgG | ln[Ag]° | -12,9132 | -13,493 | -12,3956 |
| PN13 | negative | IgG | $\ln(C_n)$ | | | |
| PN13 | negative | IgG | ν | | | |
| PN13 | negative | IgG | ln[Ag]° | | | |
| PN14 | negative | IgG | $\ln(C_n)$ | | | |
| PN14 | negative | IgG | ν | | | |
| PN14 | negative | IgG | ln[Ag]° | | | |
| PN15 | negative | IgG | $\ln(C_n)$ | | | |
| PN15 | negative | IgG | ν | | | |
| PN15 | negative | IgG | ln[Ag]° | | | |
| PN16 | negative | IgG | $\ln(C_n)$ | | | |
| PN16 | negative | IgG | ν | | | |
| PN16 | negative | IgG | ln[Ag]° | | | |
| PN17 | negative | IgG | $\ln(C_n)$ | 6,911928 | 6,606958 | 7,228209 |
| PN17 | negative | IgG | ν | 0,292047 | 0,261952 | 0,319787 |
| PN17 | negative | IgG | ln[Ag]° | -11,9653 | -12,1531 | -11,7822 |
| PN18 | negative | IgG | $\ln(C_n)$ | | | |
| PN18 | negative | IgG | ν | | | |
| PN18 | negative | IgG | ln[Ag]° | | | |
| PN19 | negative | IgG | $\ln(C_n)$ | | | |
| PN19 | negative | IgG | ν | | | |
| PN19 | negative | IgG | ln[Ag]° | | | |

| | | | |
|---|---|---|---|
| PN20 | negative | IgG | $\ln(C_n)$ |
| PN20 | negative | IgG | $\nu$ |
| PN20 | negative | IgG | ln[Ag]° |

| ID | Group | Antibody | Parameter | Value | 2.5% | 97.5% |
|---|---|---|---|---|---|---|
| PS303 | positive | IgM | $\ln(C_n)$ | 11,07182 | 10,36261 | 11,89121 |
| PS303 | positive | IgM | ν | 0,504332 | 0,481988 | 0,524805 |
| PS303 | positive | IgM | ln[Ag]° | -10,9134 | -11,3398 | -10,4352 |
| PS305 | positive | IgM | $\ln(C_n)$ | 11,5344 | 11,01921 | 12,08265 |
| PS305 | positive | IgM | ν | 0,457875 | 0,434257 | 0,479594 |
| PS305 | positive | IgM | ln[Ag]° | -11,3782 | -11,6698 | -11,0766 |
| PS308 | positive | IgM | $\ln(C_n)$ | 11,78403 | 11,11315 | 12,51167 |
| PS308 | positive | IgM | ν | 0,428395 | 0,395362 | 0,458142 |
| PS308 | positive | IgM | ln[Ag]° | -11,3626 | -11,6747 | -11,0189 |
| PS310 | positive | IgM | $\ln(C_n)$ | 11,32697 | 10,63466 | 12,06208 |
| PS310 | positive | IgM | ν | 0,426835 | 0,388578 | 0,460666 |
| PS310 | positive | IgM | ln[Ag]° | -11,7195 | -12,0404 | -11,3761 |
| PS326 | positive | IgM | $\ln(C_n)$ | 9,551911 | 8,647125 | 10,65046 |
| PS326 | positive | IgM | ν | 0,414132 | 0,373365 | 0,449884 |
| PS326 | positive | IgM | ln[Ag]° | -10,7204 | -11,227 | -10,14 |
| PS329 | positive | IgM | $\ln(C_n)$ | 11,13532 | 10,66797 | 11,62062 |
| PS329 | positive | IgM | ν | 0,296951 | 0,257886 | 0,332421 |
| PS329 | positive | IgM | ln[Ag]° | -11,9884 | -12,1772 | -11,7905 |
| PS330 | positive | IgM | $\ln(C_n)$ | 11,31267 | 10,66755 | 11,99051 |
| PS330 | positive | IgM | ν | 0,41482 | 0,376718 | 0,448585 |
| PS330 | positive | IgM | ln[Ag]° | -11,8729 | -12,1696 | -11,5603 |
| PS331 | positive | IgM | $\ln(C_n)$ | 11,88652 | 11,5773 | 12,20247 |
| PS331 | positive | IgM | ν | 0,451004 | 0,433839 | 0,467169 |
| PS331 | positive | IgM | ln[Ag]° | -11,996 | -12,1449 | -11,8439 |
| PS332 | positive | IgM | $\ln(C_n)$ | 11,35891 | 11,0365 | 11,68843 |
| PS332 | positive | IgM | ν | 0,442711 | 0,426116 | 0,458387 |
| PS332 | positive | IgM | ln[Ag]° | -12,0522 | -12,2123 | -11,8896 |
| PS352 | positive | IgM | $\ln(C_n)$ | 4,246594 | 3,513298 | 5,073963 |
| PS352 | positive | IgM | ν | 0,248676 | 0,132438 | 0,338411 |
| PS352 | positive | IgM | ln[Ag]° | -13,7019 | -15,1877 | -12,8913 |
| PS353 | positive | IgM | $\ln(C_n)$ | 11,64228 | 10,78368 | 12,58253 |
| PS353 | positive | IgM | ν | 0,435928 | 0,390541 | 0,475146 |
| PS353 | positive | IgM | ln[Ag]° | -11,5346 | -11,9589 | -11,0717 |
| PS601 | positive | IgM | $\ln(C_n)$ | 10,01357 | 9,570804 | 10,47856 |
| PS601 | positive | IgM | ν | 0,472172 | 0,452698 | 0,490287 |
| PS601 | positive | IgM | ln[Ag]° | -11,5598 | -11,8105 | -11,3024 |
| PS604 | positive | IgM | $\ln(C_n)$ | 10,61659 | 10,15826 | 11,09808 |
| PS604 | positive | IgM | ν | 0,489434 | 0,470596 | 0,50696 |
| PS604 | positive | IgM | ln[Ag]° | -11,6204 | -11,8867 | -11,3473 |
| PS607 | positive | IgM | $\ln(C_n)$ | 11,74407 | 10,97848 | 12,56237 |
| PS607 | positive | IgM | ν | 0,369802 | 0,317548 | 0,414904 |
| PS607 | positive | IgM | ln[Ag]° | -11,6626 | -11,9777 | -11,3161 |
| PS609 | positive | IgM | $\ln(C_n)$ | 6,457178 | 5,456441 | 7,548317 |
| PS609 | positive | IgM | ν | 0,359983 | 0,260567 | 0,435993 |
| PS609 | positive | IgM | ln[Ag]° | -13,2843 | -16,9079 | -12,0667 |
| PS610 | positive | IgM | $\ln(C_n)$ | 10,56408 | 9,820335 | 11,36693 |

| | | | | | | |
|---|---|---|---|---|---|---|
| PS610 | positive | IgM | ν | 0,438807 | 0,400845 | 0,472278 |
| PS610 | positive | IgM | ln[Ag]° | -11,6069 | -11,987 | -11,2015 |
| PS611 | positive | IgM | $\ln(C_n)$ | 11,0293 | 10,53715 | 11,55435 |
| PS611 | positive | IgM | ν | 0,476364 | 0,456423 | 0,494883 |
| PS611 | positive | IgM | ln[Ag]° | -11,3032 | -11,5705 | -11,0207 |
| PS619 | positive | IgM | $\ln(C_n)$ | 11,38451 | 10,60318 | 12,21617 |
| PS619 | positive | IgM | ν | 0,379671 | 0,326699 | 0,424906 |
| PS619 | positive | IgM | ln[Ag]° | -11,7287 | -12,0557 | -11,3715 |
| PS620 | positive | IgM | $\ln(C_n)$ | 11,70452 | 11,06351 | 12,38065 |
| PS620 | positive | IgM | ν | 0,416627 | 0,379945 | 0,449285 |
| PS620 | positive | IgM | ln[Ag]° | -11,76 | -12,0503 | -11,4508 |
| PS623 | positive | IgM | $\ln(C_n)$ | 11,00391 | 10,42586 | 11,61773 |
| PS623 | positive | IgM | ν | 0,476299 | 0,451714 | 0,498796 |
| PS623 | positive | IgM | ln[Ag]° | -11,6294 | -11,9435 | -11,3 |
| PS326R | positive | IgM | $\ln(C_n)$ | 10,8738 | 10,36151 | 11,41991 |
| PS326R | positive | IgM | ν | 0,481131 | 0,455966 | 0,504072 |
| PS326R | positive | IgM | ln[Ag]° | -11,2267 | -11,5016 | -10,9359 |
| PS352R | positive | IgM | $\ln(C_n)$ | 5,267515 | 4,449683 | 6,142212 |
| PS352R | positive | IgM | ν | 0,315814 | 0,208335 | 0,397578 |
| PS352R | positive | IgM | ln[Ag]° | -12,8481 | -14,047 | -12,1598 |
| PN11 | negative | IgM | $\ln(C_n)$ | 5,7918 | 5,36438 | 6,244886 |
| PN11 | negative | IgM | ν | 0,26875 | 0,224043 | 0,308585 |
| PN11 | negative | IgM | ln[Ag]° | -11,8557 | -12,1401 | -11,585 |
| PN12 | negative | IgM | $\ln(C_n)$ | | | |
| PN12 | negative | IgM | ν | | | |
| PN12 | negative | IgM | ln[Ag]° | | | |
| PN13 | negative | IgM | $\ln(C_n)$ | | | |
| PN13 | negative | IgM | ν | | | |
| PN13 | negative | IgM | ln[Ag]° | | | |
| PN14 | negative | IgM | $\ln(C_n)$ | 5,647797 | 4,38021 | 7,290554 |
| PN14 | negative | IgM | ν | 0,255935 | 0,100711 | 0,365675 |
| PN14 | negative | IgM | ln[Ag]° | -11,58 | -12,9362 | -10,5783 |
| PN15 | negative | IgM | $\ln(C_n)$ | | | |
| PN15 | negative | IgM | ν | | | |
| PN15 | negative | IgM | ln[Ag]° | | | |
| PN16 | negative | IgM | $\ln(C_n)$ | | | |
| PN16 | negative | IgM | ν | | | |
| PN16 | negative | IgM | ln[Ag]° | | | |
| PN17 | negative | IgM | $\ln(C_n)$ | | | |
| PN17 | negative | IgM | ν | | | |
| PN17 | negative | IgM | ln[Ag]° | | | |
| PN18 | negative | IgM | $\ln(C_n)$ | | | |
| PN18 | negative | IgM | ν | | | |
| PN18 | negative | IgM | ln[Ag]° | | | |
| PN19 | negative | IgM | $\ln(C_n)$ | | | |
| PN19 | negative | IgM | ν | | | |
| PN19 | negative | IgM | ln[Ag]° | | | |

| | | | | | | |
|---|---|---|---|---|---|---|
| PN20 | negative | IgM | $\ln(C_n)$ | 5,198696 | 4,289376 | 6,20635 |
| PN20 | negative | IgM | $\nu$ | 0,260222 | 0,159102 | 0,339623 |
| PN20 | negative | IgM | ln[Ag]° | -11,9612 | -13,6253 | -11,1848 |
